# TISSUE: uncertainty-calibrated prediction of single-cell spatial transcriptomics improves downstream analyses

**DOI:** 10.1101/2023.04.25.538326

**Authors:** Eric D. Sun, Rong Ma, Paloma Navarro Negredo, Anne Brunet, James Zou

## Abstract

Whole-transcriptome spatial profiling of genes at single-cell resolution remains a challenge. To address this limitation, spatial gene expression prediction methods have been developed to infer the spatial expression of unmeasured transcripts, but the quality of these predictions can vary greatly. Here we present TISSUE (Transcript Imputation with Spatial Single-cell Uncertainty Estimation) as a general framework for estimating uncertainty for spatial gene expression predictions and providing uncertainty-aware methods for downstream inference. Across eleven benchmark datasets, TISSUE provides well-calibrated prediction intervals for predicted expression values. Moreover it consistently reduces false discovery rates for differential gene expression analysis, improves clustering and visualization of predicted spatial transcriptomics, and improves the performance of supervised learning models trained on predicted gene expression profiles. Applying TISSUE to a MERFISH spatial transcriptomics dataset of the adult mouse subventricular zone, we identified subtypes within the neural stem cell lineage and developed subtype-specific regional classifiers. TISSUE is publicly available as a flexible wrapper method for existing spatial gene expression prediction methods to assist researchers with implementing uncertainty-aware analyses of spatial transcriptomics data.

## 1 Introduction

Spatial transcriptomics technologies extend high-throughput characterization of gene expression to the spatial dimension and have been used to characterize the spatial distribution of cell types and transcripts across multiple tissues and organisms [44, 9, 45, 25, 47, 64]. A major trade-off across all spatial transcriptomics technologies is between the number of genes profiled and the spatial resolution such that most spatial transcriptomics technologies with single-cell resolution are limited to the measurement of a few hundred genes but typically not the whole transcriptome [33]. Given the resource-intensive nature of single-cell spatial transcriptomics data acquisition, computational methods for upscaling the number of genes and/or predicting the expression of additional genes of interest are highly desirable.

There exist several methods for imputing or predicting spatial gene expression using a paired single-cell RNA-seq dataset. Generally, these methods proceed by joint embedding of the spatial and RNA-seq datasets and then predicting expression of new spatial genes by aggregating the nearest neighboring cells in the RNA-seq data [1, 55, 2, 66] or by joint probabilistic modeling, mapping, or transport [39, 10, 61, 64, 13, 46]. For example, SpaGE relies on joint embedding of spatial and RNAseq data using PRECISE domain adaptation followed by k-nearest neighbors regression [48, 1]; a method referred to as “Harmony” here relies on Harmony integration of the two data modalities and averaging of nearest cell expression profiles [2]; and Tangram uses an optimal transport framework with deep learning to devise a mapping between single-cell and spatial transcriptomics data [10]. Applications of these methods have been used in the characterization of spatial differences in aging of mouse neural and glial cell populations [2], recovery of immune signatures in primary tumor samples [61], and identification of spatial patterns in gene regulation [10].

Since the relative performance of these models varies significantly depending on the application area and underlying datasets, there is no best model across all use cases [33]. Moreover, variability in model performance may adversely affect downstream analysis, particularly in promoting false discoveries due to prediction errors. At the same time, few existing gene expression prediction methods provide uncertainty measures for the predicted expression profiles and there are no approaches for utilizing uncertainty in down-stream analyses like differential gene expression analysis, hypothesis testing, clustering, visualization, or supervised learning. As a result, it is often difficult to rely on predicted spatial gene expression profiles without significant external validation or understanding of their context-specific uncertainties.

Here, we present TISSUE (Transcript Imputation with Spatial Single-cell Uncertainty Estimation) as a general wrapper framework around any spatial gene expression imputation or prediction model that produces well-calibrated uncertainty measures that are tailored to the context of each individual model and its use case. We show that TISSUE can be leveraged for improvements in various uncertainty-aware data analysis tasks including the calculation of prediction intervals, hypothesis testing using a multiple imputation approach, supervised learning (e.g. cell type classification, anatomic region classification) by uncertainty-aware filtering of cells before training and prediction, and clustering and visualization using uncertainty-aware filtering and weighted principal component analysis. We further show that TISSUE can be used to identify new cell types and subtypes that have yet to be profiled using spatial transcriptomics.

## 2 Results

### 2.1 TISSUE: Cell-centric variability and calibration scores for prediction errors

Spatial gene expression prediction generally relies on leveraging spatial transcriptomics and RNAseq data from similar cell types. The RNAseq data is used to impute the expression of genes not measured in the limited spatial transcriptomics panel and can recover up to whole-transcriptome coverage of genes (Fig. 1A). To motivate the need for uncertainty quantification, we benchmarked three popular spatial gene expression prediction methods (Harmony [2], SpaGE [1], and Tangram [10]) on eight publicly available spatial transcriptomics datasets (spanning seqFISH [41], MERFISH [16], STARmap [62], ISS [29], FISH [31], osmFISH [17], and ExSeq [4] technologies; spatial data visualized in Fig. S1A) paired with single-cell or single-nuclei RNAseq datasets (spanning Smart-seq, Drop-seq, and 10X Chromium technologies) from the same organism and tissue regions [33, 38, 27, 11, 62, 23, 28, 50, 17, 4, 24, 69, 60, 54, 37, 65, 42, 71, 68]. No method consistently outperformed other methods across all spatial transcriptomics datasets. Similarly, methods that performed the best under one metric (e.g. gene-wise Spearman rank correlation between measured and predicted gene expression, see Fig. 1B) did not necessarily perform the best under an orthogonal evaluation metric (e.g. mean absolute error in predicted expression, see Fig. S1B). For a given method, there is also substantial heterogeneity in the relative performance of the model between genes and cells (Fig. 1B, Fig. S1BC), suggesting that accurate estimation of uncertainty in spatial gene expression predictions may improve confidence in interpretations and downstream analyses.

**Figure 1:**
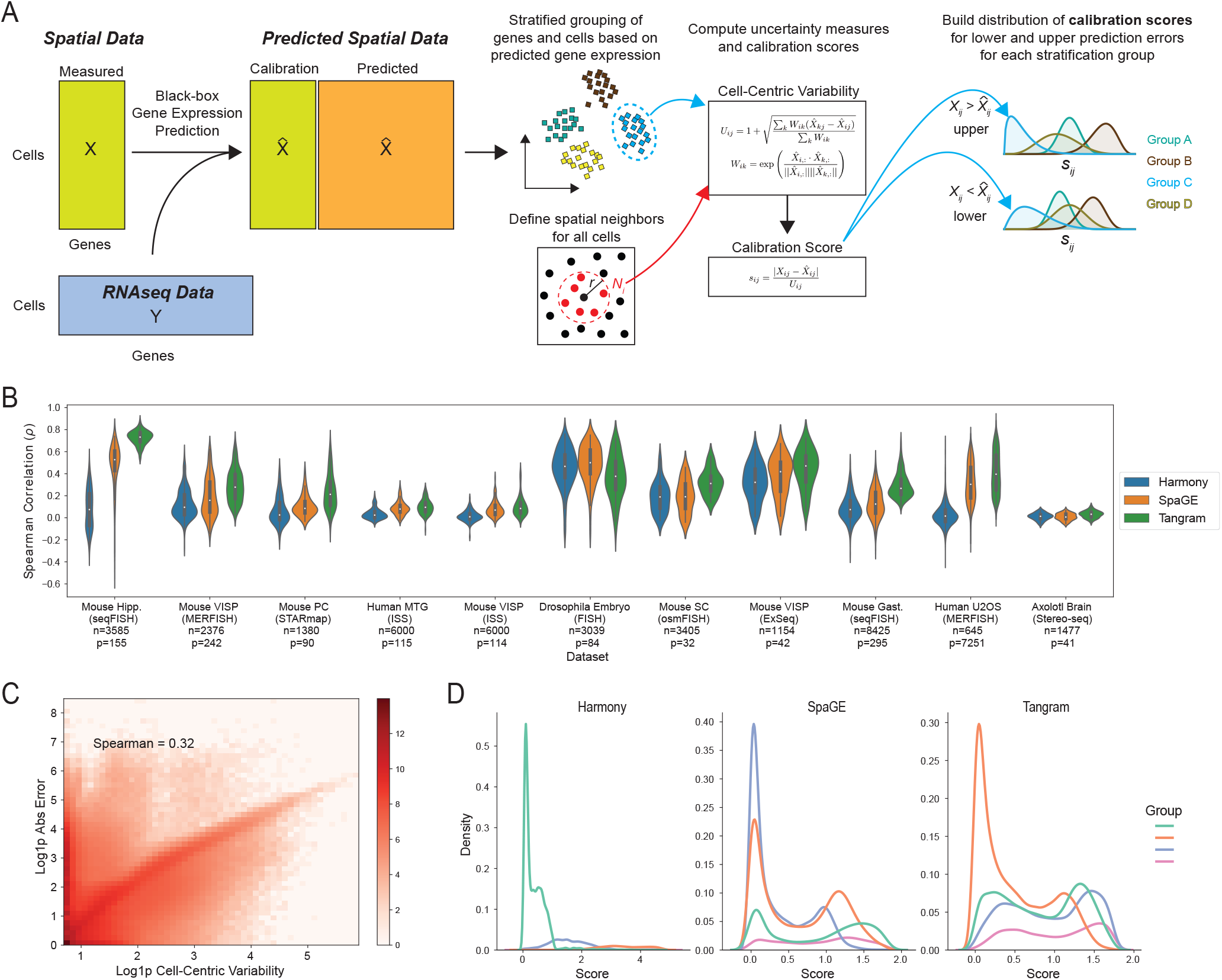
Cell-centric variability and calibration scores for conformal inference. (A) Schematic illustration of the TISSUE calibration score generation pipeline with black-box gene prediction (shown is an example method using paired spatial and RNAseq datasets), stratified grouping of genes and cells, calculation of cell-centric variability measure, and computation and allocation of the calibration score to different stratification groups. (B) Performance of three popular gene prediction methods (Harmony, SpaGE, Tangram) on eleven benchmark datasets as measured by the gene-wise Spearman correlation between predicted and actual gene expression over 10-fold cross-validation. Also shown are the number of cells (*n*) in the spatial transcriptomics datasets and the number of genes (*p*) shared between spatial and RNAseq datasets.. The inner box corresponds to quartiles of the correlation measures and the whiskers span up to 1.5 times the interquartile range of the correlation measures. (C) Correlation of TISSUE cell-centric variability and absolute prediction error across all dataset and prediction method combinations computed over 10-fold cross-validation. Log density with added pseudocount (Log1p) is shown by color, with a maximum of 1000 cells and 300 genes sampled from each dataset to provide more uniform representation. (D) Distribution of TISSUE calibration scores on mouse hippocampus ISS dataset and all three prediction method combinations using (*k*_*g*_, *k*_*c*_) = (4, 1). Details of each dataset and prediction method can be found in Methods.

Black-box machine learning models have become increasingly common across all fields of science, engineering, and medicine. In settings where there is high variability in the quality of predictions or downstream applications that require accurate predictions, it is desirable to quantify the uncertainty of model predictions. Conformal inference is a statistically rigorous and distribution-free framework for quantifying uncertainty of black-box models [7, 53, 32]. Traditionally, conformal inference proceeds by fitting a machine learning model on labeled training data, evaluating the model predictions on a small amount of labeled calibration data to build calibrated uncertainties, and then deploying the model to unlabeled test data to obtain both the predicted labels and their uncertainty. Conformal inference has been used to quantify uncertainty of region segmentations in tissue image analysis [67], measure confidence of drug discovery predictions [6], and understand the robustness of clinical treatment effects [26]. To extend the traditional conformal inference framework to spatial gene expression prediction, we make several key modifications to build well-calibrated uncertainties in TISSUE (see Methods). First, we establish an initial measure of prediction uncertainty that is scalable to unseen observations and agnostic to the prediction error. To calibrate these uncertainties to the prediction error, we build distributions of calibration scores by linking these initial measures of uncertainty to the observed prediction errors on existing genes in the spatial transcriptomics data. Finally, these calibration score distributions are used for computing well-calibrated prediction intervals and improving downstream spatial transcriptomics data analysis.

To construct an initial measure of uncertainty that can be universally applied to all existing spatial gene expression prediction methods, we posit that, on average, spatially proximate cells with similar measured gene expression profiles will also have similar expression of genes that are not measured in the spatial transcriptomics gene panel (see Fig. S2 for empirical observations supporting this assumption). As a result, large differences in predicted gene expression between neighboring cells of the same cell type would indicate low predictive performance and highly similar predicted gene expressions between neighboring cells would signify high predictive performance for the spatial gene expression prediction method. To quantify this intuition, we introduce the cell-centric variability measure, *U*_*ij*_, which, given a gene, computes for each cell a weighted measure of deviation between the predicted expression of that cell and the cells within a spatial neighborhood of it (Eq. (1) and Eq. (2)).

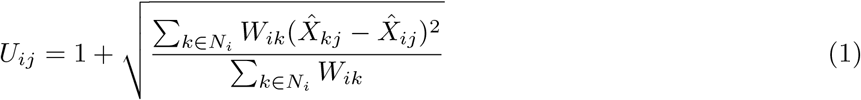

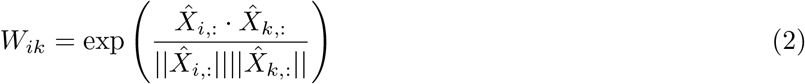

Here, 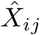 is the predicted gene expression of cell *i* and gene *j*. For a given cell *i*, its spatial neighborhood *N*_*i*_ corresponds to the *K* closest cells in the spatial transcriptomics data according to Euclidean distance. For all experiments, we use *K* = 15 but the cell-centric variability is generally robust to range of choices for *K* (Fig. S3B, see Methods for additional justifications). For each neighboring cell *k*, we compute a weight *W*_*ik*_ equal to the exponential cosine similarity in predicted gene expression profiles between the central cell *i* and its neighbor. These weights prioritize variability in predicted gene expression for similar cells (e.g. cells of the same cell type) and downplay expected variability in gene expression from dissimilar cells without the need to explicitly define cell types or states.

The cell-centric variability is generally positively correlated with the absolute prediction error for spatial gene expression (Fig. 1C). However, the cell-centric variability does not provide an exact estimate of the magnitude of these errors and the relationship between these two quantities is highly context-dependent (Fig. S3A). To explicitly link the cell-centric variability to the prediction error, we adapt a conformal inference framework by computing the calibration score, which is defined as the ratio between the absolute prediction error and the cell-centric variability (Eq. (3), Fig. 1A):

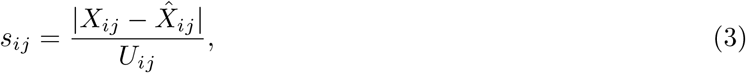

where *X*_*ij*_ denotes the measured gene expression for cell *i* and gene *j*.

The distribution of *s*_*ij*_ can subsequently be used to calibrate uncertainties for new expression predictions by multiplying the cell-centric variability of those predictions by specific quantiles of the *s*_*ij*_ calibration score set, which returns values on the scale of prediction errors (see next section for details). To permit flexible calibration schemes within the same spatial transcriptomics dataset, TISSUE allocates calibration scores to disjoint groups of genes and cells, referred to as the stratified calibration sets or groups, which are determined by k-means clustering of genes and then k-means clustering of cells by predicted gene expression (Fig. 1A, see Methods for further details). This stratified grouping scheme is motivated by the observation that there is generally positive correlation in pairwise similarities of predicted expression and of prediction error (Fig. S3C). The number of gene and cell subsets (*k*_*g*_, *k*_*c*_) can be user-specified or determined using an automated method (see Methods), but downstream results are generally robust to exact specifications of these stratified groupings. Empirically, the distribution of calibration scores can vary significantly across different identified subsets, suggesting the identification of heterogeneous calibration score sets (Fig. 1D and Fig. S3D with (*k*_*g*_, *k*_*c*_) = (4, 1)). Due to the asymmetric distribution of transcript counts, the calibration scores are further separated by the sign of the prediction error into a lower set for 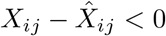 and upper set for 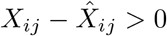 (Fig. 1A).

### 2.2 TISSUE provides prediction intervals for predicted gene expression

We leverage a conformal inference framework to convert cell-centric variability of new spatial gene expression predictions into well-calibrated prediction intervals using the calibration scores derived from the measured gene panel. Given a gene expression prediction for cell *a* and gene *b* and confidence level *α*, we compute the cell-centric variability *U*_*ab*_ and multiply it by the (⌈(*m* + 1)(1 − *α*)⌉*/m*)-th quantile of the upper and lower calibration sets of *s*_*ij*_ corresponding to all *m* predicted expression values from the same stratified group as the predicted expression of cell *a* and gene *b*, which yields the asymmetric TISSUE prediction interval with approximate 1 − *α* coverage (Fig. 2A). Using this approach, TISSUE prediction intervals can be obtained for every predicted gene expression and every cell in the spatial transcriptomics data (see Methods for details and mathematical guarantees). The TISSUE prediction interval width is positively correlated with the absolute prediction error of measured genes under cross-validation (Fig. 2B). This trend persists after normalizing by the magnitude of predicted expression (Fig. S4D) and also exists for different choices of *α* for computing prediction interval widths (Fig. S4EF). For individual genes of interest, the TISSUE prediction interval width generally reflects the spatial distributions of absolute prediction errors such as for *Plp1* in osmFISH profiling of mouse somatosensory cortex (Fig. 2C), *Neta* in a virtual spatial transcriptomics profile of *Drosophila* embryo (Fig. 2D), and *Tnfaip6* in MERFISH profiling of mouse primary visual cortex (Fig. 2E). Averaged across all genes and cells, the TISSUE prediction interval provides well-calibrated coverage of prediction errors on unseen genes for a broad range of confidence levels and across all prediction methods and spatial trancriptomics datasets (Fig. 2F). For individual genes, there is a general tendency towards well-calibrated prediction intervals (Fig. S4A). Similar calibration quality for prediction intervals was observed under automated selection of *k*_*g*_ and *k*_*c*_ (Fig. S4B). The calibration quality of TISSUE was also highly reproducible across technical replicates within a spatial transcriptomics dataset (Fig. S4C).

**Figure 2:**
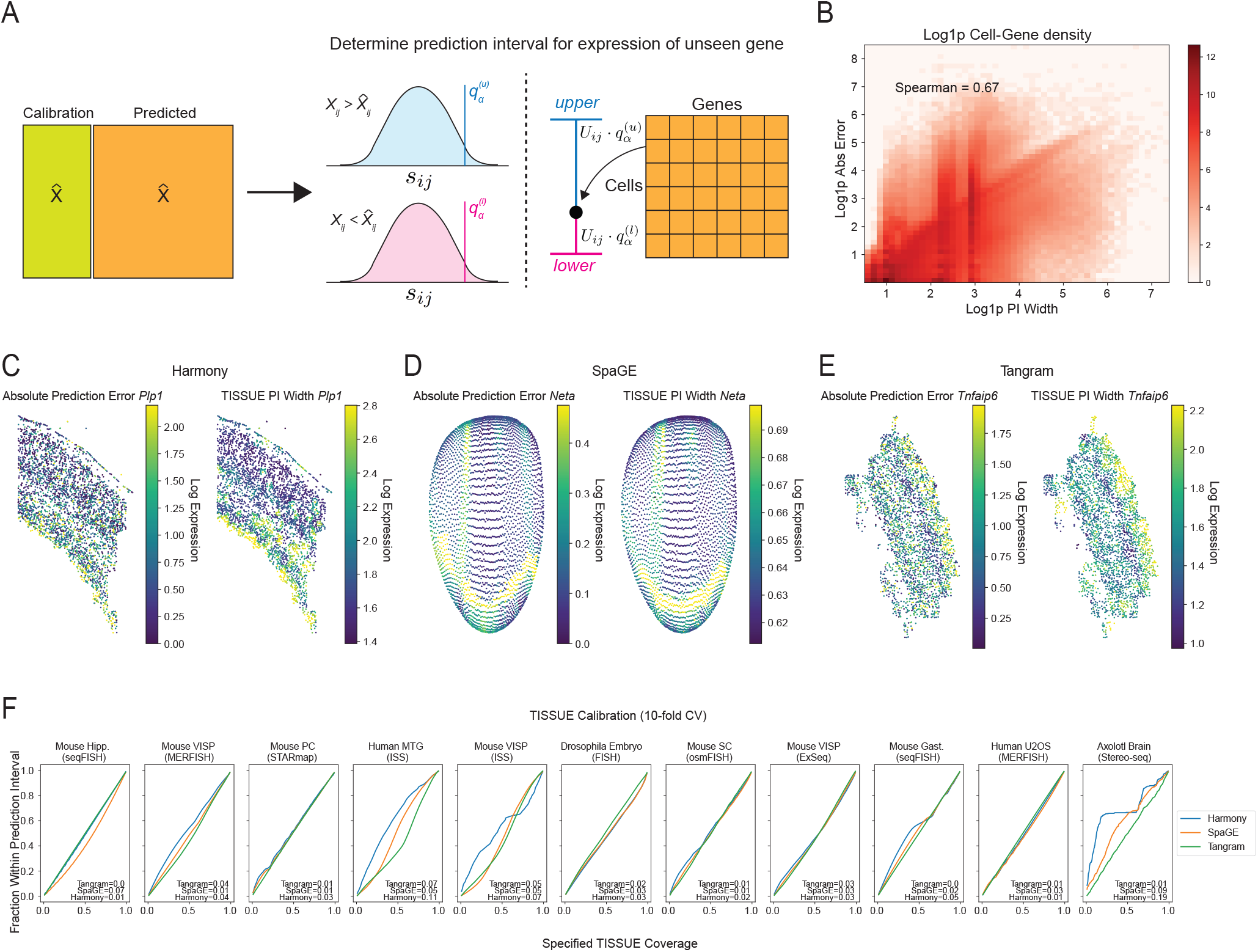
Prediction intervals for spatial gene expression. (A) Schematic illustration of the TISSUE prediction interval retrieval process from the calibration scores for a given confidence level. (B) Correlation of the 67% prediction interval width and the absolute prediction error across all dataset and prediction method combinations computed over 10-fold cross-validation. Log density with added pseudocount (Log1p) is shown by color, with a maximum of 1000 cells and 300 genes sampled from each dataset to provide more uniform representation. (C) Comparison of absolute prediction error (left) and the 67% prediction interval width (right) for a representative gene in the mouse somatosensory cortex osmFISH dataset; (D) in the virtual Drosophila embryo spatial transcriptomics dataset; (E) and in the mouse primary visual cortex MERFISH dataset. (F) Calibration curves for TISSUE prediction intervals showing empirical coverage as a function of the specified confidence level across 10-fold cross-validation. The calibration error is annotated for each prediction method (see Methods). All prediction intervals were generated with (*k*_*g*_, *k*_*c*_) = (4, 1) settings for stratified grouping.

We used Sprod, a framework for de-noising spatially resolved transcriptomics [63], and the mouse somatosensory cortex osmFISH dataset to investigate whether TISSUE calibration would be affected by spatial de-noising or alternative formulations of the TISSUE neighborhood graph for computing cell-centric variability. Across different combinations of pre-processing (with or without Sprod de-noising) and neighborhood graphs (TISSUE or Sprod cell similarity graph), TISSUE calibration quality was high and comparable to the default TISSUE settings (Fig. S4G), suggesting that the TISSUE framework is likely to be robust to de-noising and alternative cell-cell similarity graphs for defining cell neighborhoods. Similarly, the TISSUE prediction interval widths were highly correlated between the default TISSUE neighborhood graph and the alternative Sprod neighborhood graph (Fig. S4H). We also tested on gimVI, a deep generative model for spatial gene expression prediction [39], and found well-calibrated TISSUE prediction intervals for gimVI (Fig. S5).

### 2.3 Uncertainty-aware hypothesis testing and differential gene expression analysis with TISSUE

Hypothesis testing of differences in gene expression between experimental conditions, cell types, or other groupings is an important tool in understanding biological heterogeneity and perturbation effects using spatial transcriptomics. We extend TISSUE calibration scores for more robust hypothesis testing of differential predicted gene expression across conditions. Specifically, TISSUE hypothesis testing involves sampling multiple imputations for the predicted gene expression values by first sampling calibration scores 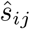’s from corresponding calibration sets and then perturbing the original predicted expression values by *U*_*ij*_ · 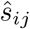 with the direction of the perturbation dependent on whether the sampled score was in the upper or lower set (Fig. 3A). Repeating this process *D* times yields *D* possible imputations for each cell and gene. Using multiple imputation theory, TISSUE then derives corrected measures of statistical significance using a modified independent two-sample t-test (Fig. 3A, see Methods for further details). These corrected statistics account for the uncertainty in prediction as encoded by the sampling of scores for generating new imputations. This multiple imputation framework can be extended to other statistics of interest [51, 3, 35, 34].

**Figure 3:**
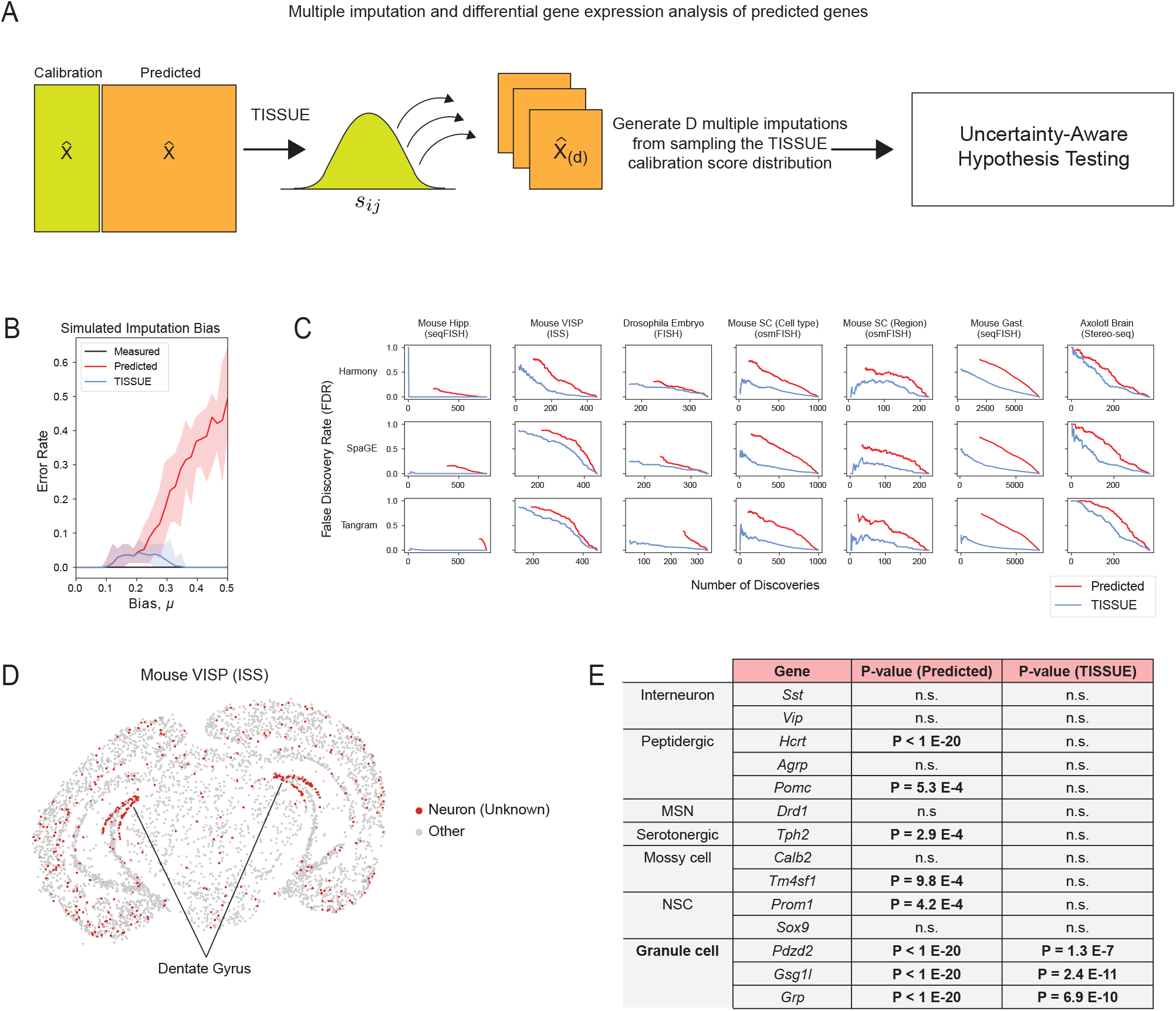
Uncertainty-aware differential gene expression analysis with TISSUE. (A) Schematic illustration of the TISSUE multiple imputation pipeline for hypothesis testing. Calibration scores are randomly sampled and used to compute new predicted gene expression profiles and statistics are compiled across all imputations using a modified two-sample t-test. (B) Error rate for testing of significant gene expression differences between two homogeneous groups of cells (*n* = 980, *p* = 1000) as a function of the selective bias in prediction error in approximately half of the genes for one group of cells using a Benjamini-Hochberg correction for 5% false discovery rate. Shown are error rates for two-sample t-test on the measured gene expression profiles (black), two-sample t-test on predicted gene expression (red), and modified two-sample t-test using the TISSUE multiple imputation approach on predicted gene expression (blue). Results were obtained using automated stratified grouping (see Methods). Bands represent the range and solid line denotes the mean error rate measured across 20 simulations. (C) False discovery rate of differentially expressed genes between cell type or anatomic region labels (one versus all approach) using the differentially expressed genes on the measured gene expression profiles as the ground truth across different p-value cutoffs. Discoveries are assessed across all genes for all class labels. Shown are results for all three prediction methods and all spatial transcriptomics datasets with cell type or region labels available. All calibration scores were generated with (*k*_*g*_, *k*_*c*_) = (4, 1) settings for stratified grouping. (D) Mouse primary visual cortex ISS dataset with the unknown neuronal cluster colored in red and spatially localized to the dentate gyrus of the hippocampus. (E) Marker genes for the unknown neuronal cell cluster are differentially expressed for multiple neuronal cell type gene sets using traditional hypothesis testing with two-sample t-test on the predicted gene expression (Predicted), but are selectively differentially expressed for granule cell marker genes using the modified two-sample t-test with multiple imputation from TISSUE calibration scores (TISSUE). P-values are shown for all predicted neuronal marker genes with significance threshold of Bonferroni-adjusted *p <* 0.05. All calibration scores were generated with (*k*_*g*_, *k*_*c*_) = (4, 1) settings for stratified grouping.

To compare TISSUE hypothesis testing to traditional hypothesis testing using only the predicted gene expression values, we generated synthetic data using SRTsim [72] in which there are two groups of cells with the same ground truth gene expression (see Methods for simulation settings). To simulate spatial gene expression prediction, we added biased Gaussian noise with mean *µ* to a portion of the genes in one of the cell groups but not the other, and standard Gaussian noise to all other gene expression values. Under this context, TISSUE hypothesis testing exhibited lower error rate in calling differentially expressed genes between the two cell groups (automated stratified grouping, Benjamini-Hochberg corrected p-value cutoff for FDR=5%) across different levels of prediction bias than traditional hypothesis testing (Fig. 3B).

In order to further evaluate TISSUE hypothesis testing, we compared it to the traditional hypothesis testing approach on seven publicly available spatial transcriptomics datasets with associated cell type or anatomic region labels (see Methods for details on data labeling). For each label, we computed statistical significance of gene expression differences within that label as compared to all cells with different labels (i.e. one-versus-all approach). Statistical significance was assessed for all genes in the measured gene expression with traditional hypothesis testing and in the predicted gene expression with both TISSUE and traditional hypothesis testing. Using the differentially expressed genes detected using measured gene expression values as the ground truth, we observed lower false discovery rate of differentially expressed genes using TISSUE hypothesis testing as compared to traditional hypothesis testing across all prediction methods and datasets. The lower FDR was observed across different numbers of differentially expressed gene detections (Fig. 3C) and across different p-value cutoffs (Fig. S6A). This decrease in FDR was also observed for automated selection of *k*_*g*_ and *k*_*c*_ (Fig. S6B). The TISSUE multiple imputation framework can also be extended to non-parametric hypothesis testing and to spatially variable gene detection with SpatialDE [59], resulting in similar reductions in false discovery rates when using predicted gene expression profiles as input, especially when the number of intended discoveries is low (Fig. S6CD). TISSUE hypothesis testing was also reproducible across different replicates within the same spatial transcriptomics dataset (Fig. S6E). As such, application of TISSUE hypothesis testing robustly guards against false discoveries when performing differential gene expression analysis with predicted spatial gene expression profiles.

To illustrate a specific use case of TISSUE hypothesis testing, we applied the method to an in situ sequencing (ISS) mouse primary visual cortex dataset. Using unbiased Leiden clustering, we identified several broad neuronal cell type clusters along with specific non-neuronal cell type clusters. We were unable to further resolve the neuronal cell clusters and used spatial gene prediction with SpaGE to predict the expression of additional neuronal subtype markers that were not in the original ISS gene panel. For one neuronal cluster, which localized to the dentate gyrus of the hippocampus (Fig. 3D) differential expression was detected across most neuronal subtype markers, but under TISSUE hypothesis testing, we observed that this cluster had selective differential expression of predicted marker genes associated with granule cells (*Pdzd2, Gsg1l, Grp*), which are concentrated in the dentate gyrus (DG) of the hippocampus [8], and no significant expression of other cell type markers, including those for mossy cells (*Calb2, Tm4sf1*) and neural stem cells (*Prom1, Sox9*), which are other cell types found in the DG [8], and those for other neuronal subtypes (*Sst, Vip, Hcrt, Agrp, Pomc, Drd1, Tph2*) (Fig. 3E). The identification of this neuronal cell cluster as a granule cell cluster was confirmed by spatial localization of these cells to the hippocampal DG and by further confirmation with measured gene marker *Lrtm4*, which has been previously implicated with granule cell processes [56]. Under traditional differential expression testing with the predicted spatial gene expression values, we were unable to recover the same specificity for granule cell markers. We observed similar but reduced trends for Tangram prediction and lack of significance for any markers under Harmony prediction, likely due to the low performance of that model on this dataset (Fig. 1B) and suggesting that comparison of TISSUE differential expression results from multiple prediction methods is advantageous.

### 2.4 Uncertainty-aware supervised learning, clustering, and visualization with TISSUE

Supervised learning is a common practice with single-cell and spatial transcriptomic data and can lead to useful models for predicting quantities of interest such as biological age [12], cell cycle state [52], and perturbational responses [40]. Similarly, cell clustering and visualization are commonly used to identify cell types in spatial transcriptomics data and intuitively understand high-dimensional differences between different groups. Substantial errors in spatial gene expression prediction may adversely affect the performance of these downstream tasks when relying on the predicted gene expression profiles as input. Here we introduce TISSUE cell filtering as an approach for retrieving a high-quality subset of predictions to be used as input for improved downstream training and evaluation of supervised classification models, clustering of cells, and data visualization via dimensionality reduction. TISSUE cell filtering involves ranking of cells by the magnitude of uncertainty (i.e. prediction interval width) for each gene, followed by automated filtering out of cells with the highest uncertainty ranking within each class label (e.g. cell type) (Fig. 4A, see Methods for details).

**Figure 4:**
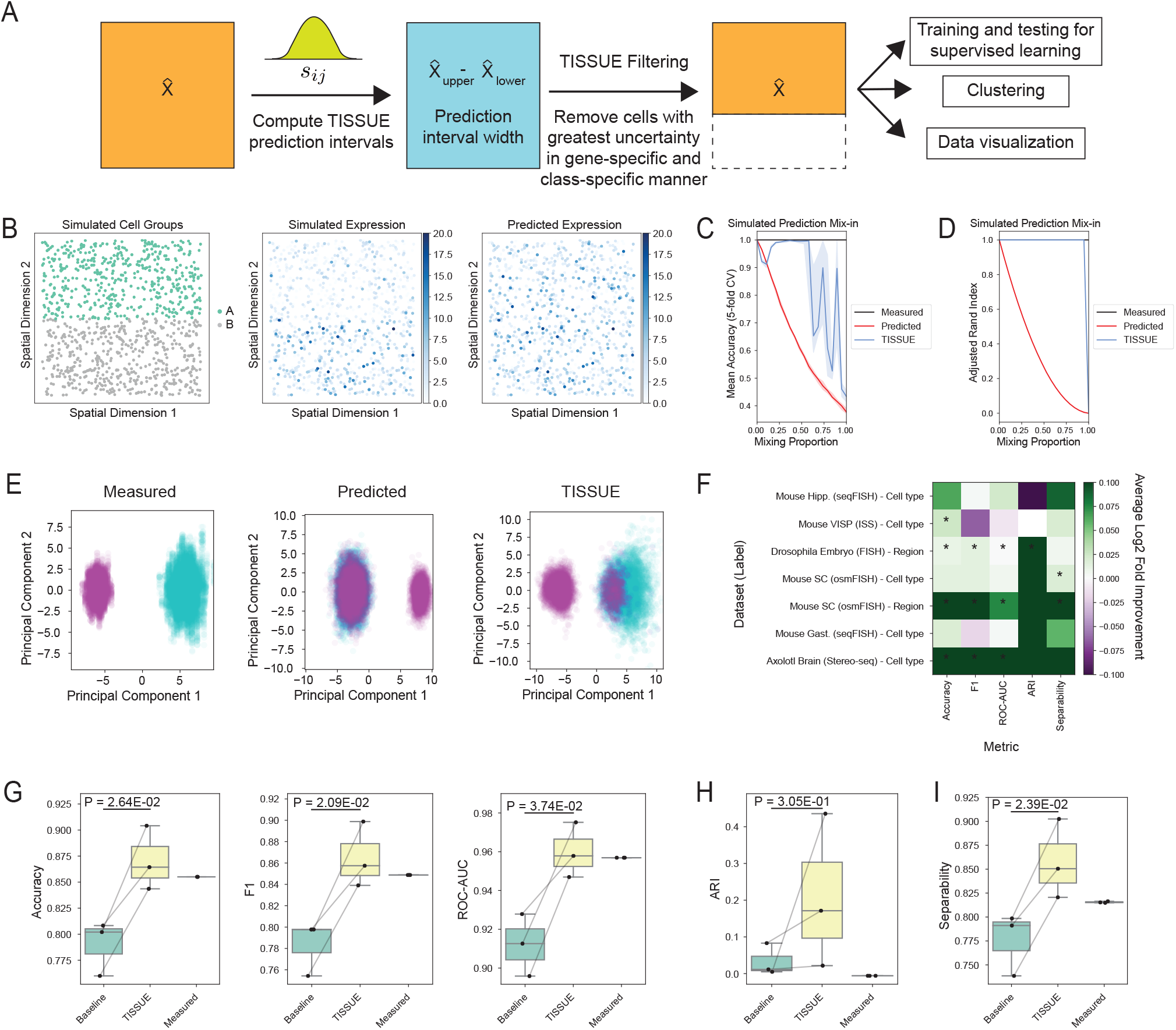
Uncertainty-aware supervised learning, clustering, and visualization. (A) Schematic illustration of the uncertainty-aware pipeline where the TISSUE prediction interval width is used to filter out cells with poor gene expression predictions from the data before downstream tasks such as supervised learning, clustering, or visualization. (B) Spatial visualization of the simulated dataset by two cell clusters (left), measured expression of a representative gene that is higher in one cluster (middle), and the predicted expression with mix-in prediction bias of the same gene (right). (C) Mean accuracy for logistic regression models trained to classify the two cell clusters from the simulated dataset as a function of the mix-in bias. Accuracy was measured across 5-fold cross-validation for models trained and evaluated on the measured gene expression values (black), predicted gene expression values (red), and TISSUE-filtered predicted gene expression values (blue). Results were obtained using automated stratified grouping. Bands represent the interquartile range and solid line denotes the median performance measured across 20 simulated datasets. (D) Adjusted rand index of k-means clustering on the top 15 principal components with *k* = 2 on the simulated dataset as a function of the mix-in bias. Colors and bands are defined similarly to panel C. (E) DynamicViz scatter plots of all cells by the first and second principal components from the measured gene expression profiles, predicted gene expression profiles, and predicted gene expression profiles after TISSUE filtering with 90% mixing of the two synthetic cell clusters. DynamicViz visualization was generated by rigidly aligning 20 PCA visualizations of different simulations (see Methods for details). Cells are colored to represent the two simulated clusters. Results were obtained using stratified grouping settings of (*k*_*g*_, *k*_*c*_) = (2, 2). (F) Log2 fold change in improvement of performance metrics using TISSUE-filtered PCA in lieu of regular PCA on predicted expression for supervised learning (logistic regression models trained to predict class labels; Accuracy, F1, ROC-AUC), clustering (k-means clustering on top 15 principal components; Adjusted Rand Index (ARI)), and visualization (linear separability measured as the classification accuracy of a linear kernel support vector classifier fitted on the top 15 principal components) for the top three classes across all dataset and class label combinations. (G-I) Downstream task performance metrics for the three most prominent anatomic region class labels for the mouse somatosensory osmFISH dataset. Shown are metrics for all three prediction methods with stratified grouping settings (*k*_*g*_, *k*_*c*_) = (4, 1). P-value was computed using a paired two-sample t-test. The box corresponds to quartiles of the metrics and the whiskers span up to 1.5 times the interquartile range of the metrics. (G) Accuracy, F1 score, and ROC-AUC (receiver-operator characteristic area under the curve) metrics for logistic regression models trained on the predicted gene expression (baseline), TISSUE-filtered predicted gene expression, or measured gene expression for classification. (H) Adjusted rand index (ARI) for k-means clustering (*k* = 3) on the top 15 principal components obtained from the predicted gene expression (baseline), TISSUE-filtered predicted gene expression, or measured gene expression for classification. (I) Linear separability measured as classification accuracy of linear kernel support vector classifier fitted on the top 15 principal components obtained from the predicted gene expression (baseline), TISSUE-filtered predicted gene expression, or measured gene expression for classification.

To compare the performance of TISSUE cell filtering to traditional approaches to downstream tasks, we generated synthetic spatial transcriptomics data using SRTsim [72] with two distinct cell type clusters where half of the profiled genes are higher in expression in one cell type (see Methods for full simulation details). Predicted gene expression is simulated by selectively adding mix-in bias to a proportion of cells in one cell type such that the expression profiles of those cells resemble the ground truth of the other cell type, and zero-centered Gaussian prediction noise is added to all other cells (see Methods for details, Fig. 4B). Under this simulation setting, supervised learning classification models trained and evaluated on TISSUE-filtered data generally outperformed classifiers trained and evaluated on unfiltered spatial transcriptomics data in separating the two cell groups across different levels of mix-in prediction bias and under cross-validation (Fig. 4C). Similarly, clustering of cells (k-means with *k* = 2 on the top 15 principal components) using the TISSUE-filtered gene expression predictions resulted in higher quality clustering than clustering of cells using the unfiltered gene expression predictions as evidenced by higher adjusted rand index (ARI) with respect to the true cell groups across different levels of mix-in prediction bias (Fig. 4D).

To assess improvements in low-dimensional visualization of the data, we used DynamicViz [58] to rigidly align cells by their top two principal components across 20 independent simulations and observed that TISSUE-filtered visualizations were better able to separate the two cell groups while the unfiltered visualizations were unable to do so under 50% mix-in of the two cell groups (Fig. 4E). The DynamicViz variance score was also lower for the TISSUE-filtered visualization than for the unfiltered PCA visualization (median variance score of 0.198 compared to 0.381), indicating more stable visualization quality in the former, likely due to the improved representation of differences between the two cell type clusters.

To assess improvements by TISSUE cell filtering on publicly available spatial transcriptomics datasets, we curated seven pairings of datasets and class labels (e.g. cell type or anatomic region) and restricted our analyses to the three labels with greatest representation within each pairing. To evaluate supervised learning for classification, we compared the cross-validated performance of logistic regression models trained and evaluated on the TISSUE-filtered predicted spatial gene expression to the performance of logistic regression models trained and evaluated on the unfiltered predicted gene expression profiles. Across three different performance metrics (accuracy, area under the receiver-operator characteristic curve, and F1 score), the TISSUE-filtered classifiers generally outperformed the unfiltered classifiers on prediction tasks (Fig. 4F), particularly for the osmFISH mouse somatosensory cortex dataset with region labels, where classification performance was comparable to that of models trained on the measured gene expression profiles (Fig. 4G). Similarly, clustering quality and visualization quality were generally improved by TISSUE cell filtering as evidenced by higher adjusted rand index (ARI) with respect to the class labels and by higher linear separability of classes in low-dimensional PCA representations of the predicted gene expression, which was measured by fitting a support vector classifier with a linear kernel to the top 15 principal components (Fig. 4FG).

For clustering and visualization tasks, we considered an alternative framework to TISSUE cell filtering, where instead of filtering out high-uncertainty cells, we leveraged weighted principal component analysis (WPCA) [18] with weights corresponding to a transformation of the inverse TISSUE prediction interval width for each cell and gene expression prediction (Fig. S7A, see Methods for further details). After applying this approach, referred to as TISSUE-WPCA, to the predicted gene expression data, we then extracted a subset of the top principal components for subsequent clustering and visualization of cells based on their gene expression. The TISSUE-WPCA approach improved linear separability between the two simulated cell groups on the synthetic datasets for a range of mix-in bias levels (Fig. S7BC) and improved clustering on several real spatial transcriptomics datasets (Fig. S7D).

### 2.5 TISSUE resolves cell types and identifies spatial heterogeneity markers of the subventricular zone neurogenic niche

The subventricular zone (SVZ) neurogenic niche is located in the lateral ventricles of the adult mammalian brain and is home to resident neural stem cells that are important in homeostasis and for injury response and repair [49, 19, 5]. In addition to many other cell types, the subventricular zone contains cells of the neural stem cell lineage, which consists of neural stem cells (quiescent and activated subtypes), neural progenitor cells, and neuroblasts, all of which have yet to be identified using spatial transcriptomics of the mammalian brain. To test the ability of TISSUE to identify and characterize these cell types for the first time in spatial transcriptomics data, we used MERFISH to profile the spatial expression of 140 genes on a young adult mouse brain section containing both lateral ventricles (Fig. 5A). We performed clustering on the data and using known marker genes, we identified several cell clusters including astrocytes, endothelial cells, ependymal cells, microglia, neurons, oligodendrocyte progenitor cells (OPCs), oligodendrocytes, and an ambiguous cell cluster that localized to the lateral ventricles (Fig. 5AB). Although the MERFISH gene panel contained several known transcriptomic markers for quiescent neural stem cells (qNSC/astrocytes), activated neural stem cells/neural progenitor cells (aNSC/NPCs), and neuroblasts (see Methods), they were insufficient for resolving the ambiguous cell cluster further into these subtypes (Fig. S9A). As such, we used SpaGE and a single-cell RNAseq dataset of the adult mouse subventricular zone [12] to predict additional genes that were not present in the original panel, including general NSC and qNSC/astrocyte markers (*Slc1a3, Nr2e1, Sox9, Vcam1, Hes5, Prom1, Thbs4*), aNSC/NPC markers (*Pclaf, H2afx, Rrm2, Insm1, Egfr, Prom1, Mcm2, Cdk1*), and neuroblast markers (*Stmn2, Dlx6os1, Igfbpl1, Sox11, Dlx1*). After sub-clustering the ambiguous cluster and then leveraging TISSUE multiple imputation and hypothesis testing, we observed differential expression of marker genes, which identified qNSC/astrocyte, aNSC/NPC, and neuroblast cell clusters (Fig. 5C). In particular TISSUE multiple imputation and hypothesis testing was necessary to resolve the identity of the aNSC/NPC subcluster, which could not be resolved from the SpaGE-imputed expression values alone (Fig. S9B). Consistent with known biology of the SVZ, all three cell subtypes were found throughout both lateral ventricles (Fig. 5D) and the relative proportions of each subtype was similar between the right and left ventricles (Fig. S9C). Additionally, there were slightly more neuroblasts than aNSC/NPCs in the MERFISH dataset, which is also reflected among several independent single-cell RNAseq datasets of the adult mouse subventricular zone (Fig. S9D) [21, 36, 12].

**Figure 5:**
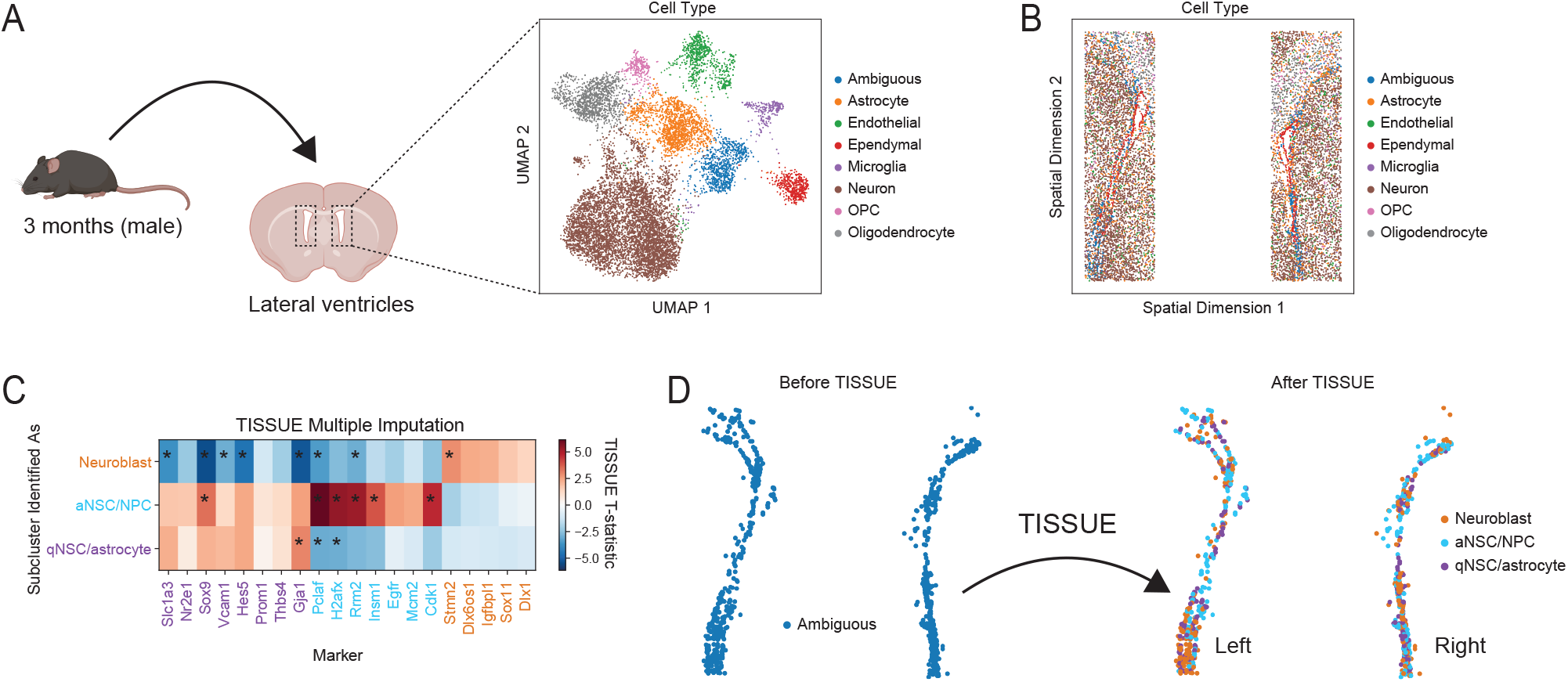
TISSUE discovers subtypes in neural stem cell lineage of the subventricular zone. (A) Schematic illustration of the 140-gene MERFISH dataset generated on adult mouse subventricular zone samples from both lateral ventricles (left) with UMAP visualization of cells colored by identified cell type clusters including one ambiguous cell cluster (right). (B) Spatial visualization of cells colored by identified cell type clusters including one ambiguous cell cluster. (C) Heatmap of TISSUE multiple imputation two-sample t-statistic values for the SpaGE predicted expression of new marker genes within cells of the three ambiguous cell sub-clusters compared to all other cells in the ambiguous cell cluster. Red boxes correspond to over-expression of that marker in that sub-cluster while blue boxes correspond to under-expression of that marker in that sub-cluster. Boxes with asterisks denote expression differences with Bonferroni-adjusted *P <* 0.05 with the TISSUE multiple imputation two-sample t-test. Colored text correspond to the identified cell subtypes and their markers, with some qNSC/astrocyte markers also known to be expressed in aNSC/NPCs. (D) Spatial visualization of the ambiguous cell type cluster before the application of TISSUE (left) and then of the three TISSUE-identified subtypes of the ambiguous cell cluster including neuroblasts, activated neural stem cells (aNSC/NPCs), and quiescent neural stem cells (qNSC/astrocytes) (right).

Recent efforts have uncovered biological heterogeneity in neural stem cell populations between the dorsal and ventral regions of the subventricular zone [14, 15]. However, these existing transcriptomic characterizations of the subventricular zone have relied on dissociated single-cell transcriptomic data, thus precluding analyses involving the ground truth spatial location of the neural stem cells without resource-intensive regional micro-dissections. Using the spatial locations of cells determined via MERFISH imaging, we categorized cells into dorsal, ventral, or other regional classes using horizontal boundaries (Fig. S9E). Using TISSUE multiple imputation and hypothesis testing, we then performed whole-transcriptome differential gene expression on dorsal or ventral categories within each of the three subtypes. For each cell subtype, we selected the 20 most differentially expressed genes and trained penalized logistic regression models to predict dorsal or ventral regional origin from predicted spatial gene expression with or without TISSUE cell filtering. In all cell subtypes, the TISSUE-filtered models outperformed the baseline unfiltered models across several classification metrics including F1 score, accuracy, area under the receiver-operator curve, and average precision (Fig. S9F). Application of these TISSUE-filtered dorsal/ventral region classifiers to dissociated single-cell RNAseq data may provide a useful first-step estimation of the regional origin of cells in the neural stem cell lineage without the need for laborious regional micro-dissections of the niche.

## 3 Discussion

Spatially resolved transcriptomics at single-cell resolution have paved the way to greater understanding of spatial patterns of gene expression and their underlying biological processes, but are currently limited to a relatively small number of detectable genes. Methods for predicting additional spatial gene expression profiles (e.g. from paired single-cell RNA-seq data) have been proposed to address this limitation. However, these methods can exhibit significant heterogeneity in prediction quality for different cells and different genes, and this heterogeneity manifests differently depending on the selected method and application. The lack of confidence measures for these predictions likely limits the adoption of these methods in spatial transcriptomics data analysis. Here, we developed a computational framework, TISSUE, to compute well-calibrated and context-specific measures of uncertainty for predicted spatial gene expression profiles.

In addition to well-calibrated prediction intervals, TISSUE provides general frameworks for leveraging uncertainty in downstream analysis such as differential gene expression testing, clustering and visualization, and supervised learning. These frameworks are flexible and can be easily adapted into existing spatial transcriptomics data analysis workflows in place of traditional analysis methods that do not account for uncertainty. For example, the differential gene expression analysis testing approach can be adapted to other hypothesis tests, which we show for non-parametric two-sample tests and spatially variable gene detection (Fig. S6CD). Likewise, the principal components obtained from TISSUE cell filtering can be used for any downstream algorithms that utilize the reduced dimensionality representation of PCA as input. Finally, the TISSUE-based filtering of training and evaluation data for supervised learning strictly modifies the data in a model-agnostic manner and therefore can be extended to both training and deployment of any supervised learning model across both regression and classification tasks. TISSUE motivates the future development and benchmarking of uncertainty-aware versions of single-cell and spatial transcriptomics data analysis methods, such as for spatial domain detection [20], embedding for label transfer [57], and cross-modality transformation [22].

As a case study, we applied TISSUE to predict the expression of additional cell type markers in the adult mouse subventricular zone MERFISH dataset. TISSUE multiple imputation and hypothesis testing were then used to successfully annotate ambiguous subtype clusters corresponding to cells in the neural stem cell lineage, which to our knowledge, constitute the first identification of these cell subtypes in spatial transcriptomics. In both the subventricular zone MERFISH dataset and in the primary visual cortex ISS dataset, TISSUE was necessary to uniquely identify ambiguous cell clusters, which was not possible using only the predicted gene expression. Together, these results indicate that TISSUE may serve as a promising framework for identifying new or previously unprofiled cell types in spatial transcriptomics data when the measured gene panel is insufficient.

TISSUE may have limited performance in contexts where spatial gene prediction patterns are not represented in the calibration set and for genes with extremely sparse expression patterns. Due to the assumptions underlying TISSUE, there may also be reduced performance on rare cell types that are not spatially co-localized. Since TISSUE performance is dependent on the size and diversity of the calibration sets, the method will generally scale better to spatial transcriptomics datasets with a large number of cells, a large number of genes, or more uniform representation of cell types. Although, specification of *k*_*g*_ and *k*_*c*_ to the natural dimensions of the data based on domain knowledge may provide optimal TISSUE performance, this performance is generally robust to most reasonable choices of *k*_*g*_ and *k*_*c*_. TISSUE is also capable of automated stratified grouping. The spatial nature of TISSUE statistics (Fig. 2C-E) may lend to spatial interpretations of the data and gene expression prediction methods, much like saliency maps have achieved for neural network models [70].

The main computational burden imposed by TISSUE is the cross-validated prediction of gene expression on the calibration set, which is necessary for building context-specific uncertainties. For *k*-fold crossvalidation with a given prediction method, TISSUE prediction would take *k* times longer than a single prediction. Alternative approaches for measuring uncertainty such as variance in predicted expression values over multiple predictions would require significantly more computation than for TISSUE cell-centric variablity to generate predictions for context-specific calibration (i.e. *k* × *l* times longer than a single prediction for the variance over *l* predictions). Here we used 10-fold cross-validation in our experiments, but in practice, the number of cross-validation folds can be specified by the user to achieve a desired runtime. The computational burden for computing cell-centric variability, calibration scores, and prediction intervals is comparatively less than that for generating the initial predictions (Fig. S10).

Although TISSUE has thus far been tested in the spatial transcriptomics setting, the underlying assumptions can generalize to other spatial data modalities, such as spatial proteomics. As whole-transcriptome spatial gene expression profiling becomes possible in the future and other spatial -omics technologies mature, we anticipate that TISSUE will find additional use in the prediction and quantification of uncertainty for enhanced spatial data analysis across multiple modalities.

## 5 Methods

### 5.1 Preprocessing of datasets

We followed data preprocessing approaches from prior benchmark comparisons of spatial gene prediction methods [33], which found highest predictive performance for these methods when non-normalized single-cell spatial transcriptomics data was paired with normalized single-cell RNAseq data. The RNAseq data was normalized using the Scanpy function pp.normalize_total() with its default settings followed by log transformation with an added pseudocount. We selected only genes expressed in at least one percent of cells in the RNAseq data.

### 5.2 Prediction of spatial gene expression

#### 5.2.1 General framework for spatial gene expression prediction

The spatial gene prediction problem involves paired data from spatial transcriptomics and RNA-seq that are approximately from the same tissue and organism. We denote the spatial transcriptomics data as *X* ∈ ℝ^*n*×*p*^ and the RNA-seq data as *Y* ∈ ℝ^*m*×*q*^, where rows are cells and columns are genes. Generally, spatial gene prediction considers the case where *q >> p* and the genes present in *X* are a subset of those in *Y* . A spatial gene prediction method predicts the expression of a gene that is present in *Y* but not in *X* for each cell in *X* using information from both *X* and *Y* .

#### 5.2.2 Harmony

Harmony (as referred to in this work) involves joint embedding of the spatial and RNAseq data using the Harmony algorithm [30] followed by k-nearest-neighbor averaging to calculate predicted expression values for each spatial cell based on its nearest neighbors in the RNAseq data. We implemented the Harmony algorithm following the description outlined in previous applications [2]. We used default Harmony settings in the Scanpy external.pp.harmony_integrate() implementation. We averaged across the 10 nearest RNAseq neighbors for each spatial cell using the first min{30, *p*} Harmony principal components.

#### 5.2.3 SpaGE

SpaGE performs spatial gene prediction using a two-step approach consisting of alignment using the domain adaptation algorithm PRECISE [48] and then performing k-nearest-neighbor regression [1]. We used a local download of the SpaGE algorithm available at https://github.com/tabdelaal/SpaGE with version corresponding to a download date of July 19, 2022. We set the number of principal vectors in SpaGE equal to 20 if *p <* 40 and to min{*n, p*}*/*2 rounded to the nearest integer if *p* ≥ 40 and otherwise used the default settings.

#### 5.2.4 Tangram

Tangram uses a deep learning framework to create a mapping for projecting RNAseq gene expression onto space [10]. We followed preprocessing details for Tangram according to previous benchmarks [33], which consisted of Leiden clustering on the scaled highly variable genes in the spatial data using Scanpy methods with default settings unless otherwise specified: pp.highly_variable_genes(), pp.scale() with max_value=10, tl.pca(), pp.neighbors(), and tl.leiden() with resolution=0.5. After preprocessing, the identified clusters were used by Tangram to project the RNAseq cells onto space using map_cells_to_space() with mode=‘clusters’ and density_prior=‘rna_count_based’ and project_genes() with default settings.

### 5.3 Calibration scores for spatial gene expression prediction

#### 5.3.1 Modifications to standard conformal inference framework

Conformal inference provides a framework for developing rigorous prediction intervals for predictions made from machine learning models. We extend this framework to construct prediction intervals for predicted gene expression values in spatial transcriptomics. To adapt this framework, we make the following key modifications and additions to the traditional theory: (1) defining a scalar measure of uncertainty (cell-centric variability) that utilizes spatial context and can be measured in a single pass of any spatial gene expression method; (2) translating from the supervised learning setting to the unsupervised setting for spatial gene expression prediction, which includes using the entire spatial transcriptomics data for calibration; (3) calculating fine-resolution prediction intervals at the level of cell-gene combinations instead of general uncertainties for a given gene or a given cell; (4) calculating asymmetric prediction intervals that are more suitable to RNA count data; (5) building custom calibration of uncertainties with hierarchically stratified groupings consisting of combinations of genes and cells; (6) designing uncertainty-aware methods and algorithms for using TISSUE prediction intervals and uncertainties for a variety of downstream data analysis tasks.

#### 5.3.2 Cross-validated spatial gene expression prediction

In order to compute calibration scores, we obtain estimated gene expression predictions on genes that are already measured in the spatial transcriptomics data. This is achieved through a cross-validation approach where a subset of the genes in the spatial transcriptomics data are left out and the gene prediction method is fitted to the remaining genes (i.e. calibration genes) to make predictions on the left-out subset of genes. In practice, we use 10-fold cross-validation to obtain predictions for all genes in the spatial transcriptomics data but the TISSUE implementation provides options to customize the cross-validation procedure according to user specifications.

#### 5.3.3 Cell-centric variability

We outline three desiderata to guide the development of a scalar uncertainty measure for spatial gene prediction:

1. To ensure computational scalability, the measure can be calculated on a single set of predicted gene expression values.
2. To accurately measure heterogeneity in prediction performance, the measure provides specific values for any cell and gene.
3. The measure ideally leverages spatial and gene expression similarity information.

We introduce the cell-centric variability to satisfy these desiderata. Specifically, for a cell *i* and gene *j*, the cell-centric variability *U*_*ij*_ is computed according to Eq. (1) and Eq. (2) using cells in its local neighborhood *N*_*i*_. We defined cell neighborhoods as the 15 nearest cells by Euclidean distance for each cell, and removed outliers by subsequently excluding neighbors with distance greater than *Q*3 + 1.5 ×(*Q*3 − *Q*1), where *Q*3 and *Q*1 are the third and first quartiles of neighbor distances across all cell neighborhoods. We used this approach for defining cell neighborhoods for all experiments since it did not require gene expression information and thus could be generalized to unseen genes; was approximately spatially scale-invariant such that in all eleven spatial transcriptomics datasets, 15 neighbors was between the average 1-hop and 2-hop neighborhood size for a Delaunay triangulation mesh graph of the cells; ensured a sufficiently high number of cells to compute the cell-centric variability reliably; and ensured a relatively fixed number of cells in each cell neighborhood such that cell-centric variability estimates could be comparable across different cells and different contexts.

The intercept term in Eq. (1) is included to ensure well-defined calibration scores and non-zero prediction interval widths for cells with no differences in gene expression across its neighborhood, which can result from the high sparsity of single-cell transcriptomics data. The weights *W*_*ik*_ in Eq. (2) are used to impose a soft weighting of the cell-centric variability for similar neighboring cells (i.e. of the same cell type) over dissimilar neighboring cells (i.e. of different cell types) without the need for user specification of discrete cell type clusters. The cell-centric variability can be computed for all cells and genes in both the calibration set (genes in the spatial transcriptomics data) and evaluation and test sets (genes to be predicted that are not in the spatial transcriptomics data).

#### 5.3.4 Calculation of calibration scores from variability measure

To link the cell-centric variability to the prediction error, we compute the calibration score as shown in Eq. (3). We compute calibration scores for all pairs of cells and genes in the calibration set (i.e. present in spatial transcriptomics data) and allocate them to their corresponding stratification groups (see following section for details). Calibration scores are further separated by the sign of 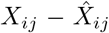 to construct non-symmetric uncertainty bounds around the predicted expression value with 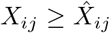 designating inclusion in the calibration scores set for the upper interval and 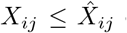 designating inclusion in the calibration scores set for the lower interval.

#### 5.3.5 Stratified cell and gene grouping for calibration scores

In addition to context-specific construction of calibration scores, TISSUE can also provide finer groupings of genes and cells, each with their own calibration score sets. These stratified groupings are specified by the number of gene groups *k*_*g*_ and the number of cell groups *k*_*c*_ for a total of *k*_*g*_ × *k*_*c*_ groups. Stratified grouping is performed for genes first through *k*-means clustering with *k* = *k*_*g*_ of the genes on the first 15 principal components representing the transposed predicted gene expression matrix. Then, within each of the identified gene strata, we perform further *k*-means clustering with *k* = *k*_*c*_ of the cells on the first 15 principal components representing the predicted gene expression matrix restricted to genes present in that strata.

Since there is no guarantee that all stratified groups will contain genes in the calibration set or that all stratified groups will have adequate number of scores for calibration, for stratified groups with less than 100 scores in either the upper interval or lower interval calibration score sets, we defaulted the calibration score set to the union of all calibration score sets across all stratified groups. To assess how representative the calibration set is for each stratified group, TISSUE includes options to measure the Wasserstein distance between the cell-centric variability of a group and that of the subset of genes in its calibration set.

TISSUE also includes an option (‘auto’) for automated selection of the stratified grouping parameters. The automated selection is achieved by first performing an increasing line search of integer *k*_*g*_ *>* 1 values and then performing *k*-means clustering of cells on the transposed predicted gene expression matrix. Finally, we compute the silhouette score on the identified clusters and increment *k*_*g*_ until the silhouette score decreases and then set *k*_*g*_ equal to the value at which maximum silhouette score was achieved. Then, we perform a similar incremental line search for *k*_*c*_ *>* 1, use *k*-means clustering of genes on the predicted gene expression matrix, and return the *k*_*c*_ value for maximum silhouette score after the same early stopping as described previously.

### 5.4 Conformal prediction intervals

#### 5.4.1 Retrieval of prediction intervals from calibration scores

For a given confidence level *α*, we construct the prediction interval with approximate probability coverage (1 − *α*) by retrieving the 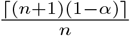-th quantiles of the upper interval calibration scores and lower interval calibration scores. Referring to these quantiles as 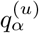 and 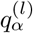 respectively, the non-symmetric conformal prediction interval for the predicted gene expression of cell *i* and gene *j* can be computed as:

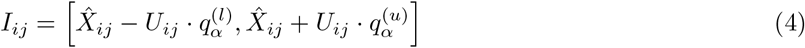

Since this prediction interval does not explicitly depend on the measured prediction error, it can be calculated for all predicted gene expression values, even if the gene was not originally present in the calibration set.

#### 5.4.2 Coverage guarantee for TISSUE prediction intervals

Under regularity conditions, the conformal inference framework provides consistent symmetric prediction intervals when applied to a scalar uncertainty measures such as cell-centric variability [7]. Building on that result, we show that this consistency is still valid with the non-symmetric prediction intervals that we compute using TISSUE.

##### Proposition 1

*Let* 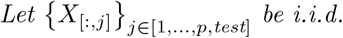 *from some distribution, then P* (*X*_*ij*_ ∈ [*l*_*ij*_, *u*_*ij*_]) ≥ 1 − *α for any confidence level* 0 ≤ *α* ≤ 1, *where l*_*ij*_ *and u*_*ij*_ *are the quantiles of the lower and upper calibration score sets corresponding to X*_*ij*_.

Here, ‘test’ refers to the index or set of indices for predicted genes that are not in the measured spatial transcriptomics data. Using the notation 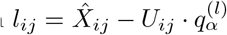 and 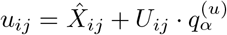, the coverage of the TISSUE prediction interval for some confidence level *α* can be represented as follows:

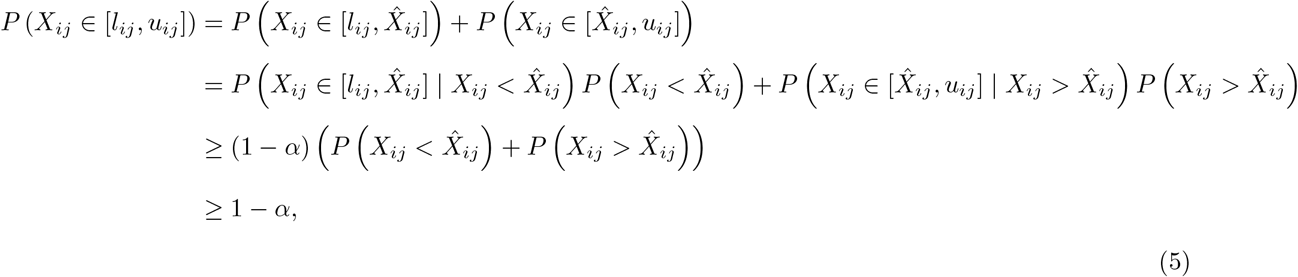

with the first inequality following from Theorem 1 of [7]. And thus, given that symmetric intervals provide proper coverage [7], then we are also guaranteed proper coverage with the asymmetric prediction intervals produced by TISSUE, which is further evident through the empirical coverage assessment for TISSUE (Fig. 2F). This guarantee extends to the stratified group setting for *k*_*g*_ *>* 1 and/or *k*_*c*_ *>* 1 (see Proposition 2 in [7]).

#### 5.4.3 Evaluation of prediction intervals

We evaluate the empirical coverage of the prediction intervals using 10-fold cross-validation splits of the genes in the spatial transcriptomics data into a calibration set and an evaluation set. We leave out the evaluation set and use the calibration set to compute calibration scores. Then using these calibration scores, for every value of *α*, we compute TISSUE prediction intervals for all cells and genes in the evaluation set. We then compute empirical coverage of the TISSUE prediction intervals, which is defined as the fraction of measured gene expression values in the evaluation set that fall within their respective TISSUE prediction interval across all cells and genes (as shown in Fig. 2F and Fig. S4B) or across all cells for each individual gene with nonzero predicted and measured expression (as shown in Fig. S4A). Well-calibrated coverage of TISSUE prediction intervals are indicated by close equivalence of the empirical coverage (“Fraction Within Prediction Interval”) and the theoretical coverage (“Specified TISSUE coverage”). For datasets with a small number of cells, there will likely be worse calibration for choices of *α* that are very close to either 0 or 1 due to sparse calibration sets for those values.

We measured the “calibration error” by measuring the area between the difference of the empirically calibration curve and the theoretically optimal calibration curve. Numerically, this involved computing the absolute difference between the empirical coverage and the theoretical coverage at each alpha value and then estimating the absolute area under this difference curve using the trapezoidal rule with default implementation of numpy.trapz().

#### 5.4.4 Sprod de-noising and alternative cell similarity graph

To investigate the effect of de-noising on TISSUE calibration performance, we used Sprod to pre-process the mouse somatosensory cortex osmFISH dataset to yield a “de-noised” version and this was used in place of the original dataset in TISSUE calibration. We used the pseudo-image option in Sprod since the corresponding images for the dataset were not publicly available. All TISSUE settings were kept identical to the settings for the original analyses.

To investigate the effect of other cell similarity graphs on TISSUE calibration, we used the cell similarity graph constructed by Sprod in the output file “sprod_Detected_graph.txt” in place of the default cosine similarity graph constructed by TISSUE. To ensure connectivity, we added the identity matrix to the Sprod cell similarity adjacency matrix, and otherwise kept all TISSUE settings at their default settings.

### 5.5 Uncertainty-aware hypothesis testing for predicted spatial gene expression

#### 5.5.1 Generating multiple imputations using calibration scores

We introduce a multiple imputation procedure for generating uncertainty-aware hypothesis testing for predicted spatial gene expression. Multiple imputations are generated by uniformly sampling calibration scores from the corresponding union of upper and lower interval calibration score sets for each predicted spatial gene expression value. Given a uniform random sample of such calibration scores, *S*_(*d*)_ ∈ ℝ^*n*×*p*^, we compute an alternative imputation as:

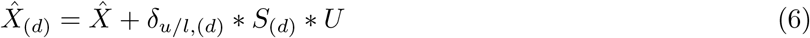

where ∗ denotes element-wise multiplication, *U* ∈ ℝ^*n*×*p*^ is equal to the cell-centric variability measures computed on 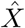, and *δ*_*u/l*_ = 1 if the score was sampled from the upper interval set and *δ*_*u/l*_ = −1 if the score was sampled from the lower interval set. This sampling is repeated *D* − 1 times to generate *D* multiple imputations including the original imputation 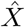. We tempered the multiple imputations against outliers by restricting the sampling to the scores within the set corresponding to the 80% conformal prediction interval.

#### 5.5.2 Modified two-sample t-test for multiple imputation

After generating *D* multiple imputations from the calibration scores, we perform hypothesis testing using a modified two-sample t-test. Consider two groups with sets of sample/cell indices *A* and *B*. Then, the mean difference and variance under normal two-sample t-test for a single imputation are:

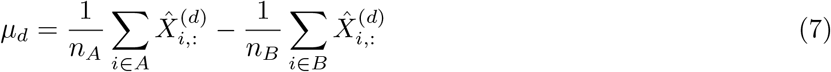

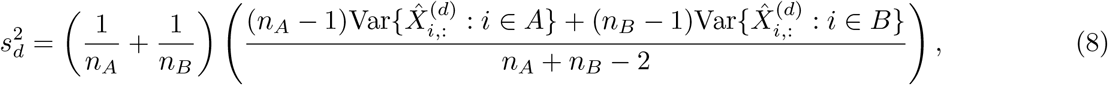

where 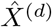 denotes the *d*-th imputation among the *D* multiple imputations. Extending these statistical measures to the multiple imputation case, we use the standard modification for multiple imputation [51, 43, 3, 35], which results in the following mean and variance:

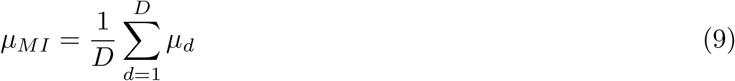

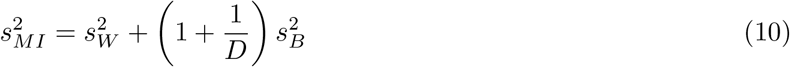

where 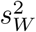 is the within-imputation variance and 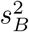 is the between-imputation variance, computed as:

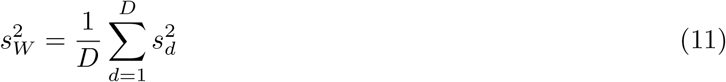

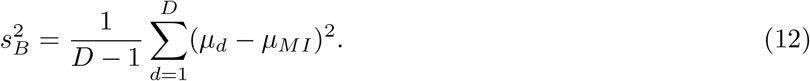

Then, the modified test statistics for the two-sample t-test is:

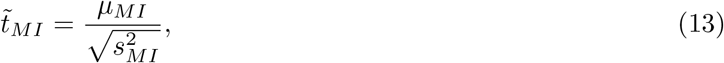

which is t-distributed with degrees of freedom 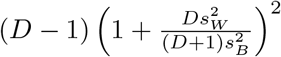 and the resulting probability can be interpreted as the posterior probability of a significant difference in means between the two groups (i.e. *β ≠* 0) accounting for both evidence of this effect and the reliability of the imputations by inflating the standard error for this effect [51]. Under some regularity assumptions (approximate normality of imputed values and missing at random), the multiple imputation approach produces consistent estimates [3, 35].

#### 5.5.3 Empirical evaluation of multiple imputation hypothesis testing

For each dataset, TISSUE hypothesis tests were computed using 10-fold cross-validation. In each cross-validation fold, we generated TISSUE spatial gene expression predictions and calibration scores on the calibration genes. Then, statistics for the TISSUE hypothesis test were computed for the held-out genes. This procedure is repeated across all folds to accrue statistics for all genes in the dataset. For TISSUE multiple imputation t-test and Wilcoxon test frameworks, we used 100 multiple imputations in each hypothesis test. For TISSUE multiple imputation SpatialDE framework, we used 10 multiple imputations in each hypothesis test due to the longer computational run-time for SpatialDE.

#### 5.5.4 Alternative multiple imputation hypothesis tests

To extend the multiple imputation testing framework to non-parametric two-sample tests, TISSUE can perform one-sided Wilcoxon/Mann-Whitney tests for ‘greater than’ or ‘less than’ comparisons between two multiple imputed samples. For these test, we use the scipy.stats.mannwhitneyu() implementation with either alternative=‘greater’ or alternative=‘less’ options. In a similar way, TISSUE can be extended to spatially variable gene detection methods such as SpatialDE [59], which provide p-values as a measure of statistical significance of spatially variable expression. We use the standard implementation and transform the input to log normalized counts before running SpatialDE.

The implementation of these frameworks is identical to the TISSUE multiple imputation t-test with the exception of using a different rule in combining inference across multiple imputations [34]. In this approach, we obtain p-values of independent hypothesis tests (i.e. one-sided Mann-Whitney test between two samples or SpatialDE across all cells with spatial coordinates) on each of *m* imputations, {*p*_1_, …, *p*_*m*_}. Then, we transform the p-values to approximate normality, {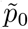, …, 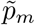}, and combine these transformed values with 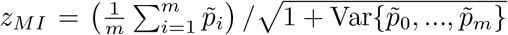. Finally, we transform the combined value back to the original scale to obtain an multiply imputed p-value estimate, *p*_*MI*_ [34]. The transformation and inverse transformation are achieved in practice with scipy.stats.norm.ppf() and scipy.stats.norm.cdf() respectively. Notably, this alternative approach involves running *m* independent hypothesis tests and is less computationally efficient than the multiple imputation t-test, which performs an aggregate test at the end of the procedure.

Due to computational constraints, we were only able to evaluate the Mann-Whitney/Wilcoxon framework on four of the seven labeled datasets, and we were only able to evaluate the SpatialDE framework on two small datasets, which were unlabeled since SpatialDE uses spatial coordinates of the cells for testing.

### 5.6 Simulated prediction bias for differential gene expression analysis

We generated synthetic data using SRTsim [72] for comparing the TISSUE uncertainty-aware hypothesis testing approach against traditional hypothesis testing. We used the reference-free SRTsim framework and generated a synthetic counts matrix using their native Shiny app. The data consists of two cell groups, referred to as ‘A’ and ‘B’, which were determined by manual drawing of a linear boundary between two spatial domains in the SRTsim Shiny application. In this setup, the separation is artificial with no simulated expression differences between the two groups. There are 465 ‘A’ cells and 515 ‘B’ cells for a total of 980 cells. To generate counts, we followed the default SRTsim recommendations and used the zero-inflated negative binomial model and set the zero proportion to 0.05, dispersion to 0.5, and mean to 2. We simulated 1000 genes, where there was no systematic difference in expression of any gene between cell group ‘A’ and cell group ‘B’. We used a random seed of 444 for SRTsim.

To simulate prediction bias for cells under condition ‘B’, we added shifted Gaussian noise (mean equal to *µ* ≥ 0, variance equal to one) to half of the genes for all cells in condition ‘B’. Standard Gaussian noise was added to the other simulated expression values (i.e. all cells and genes in group ‘A’ and the other half of genes for group ‘B’). This simulation results in prediction errors that artificially produce a difference in predicted expression between the two groups for half of the genes despite the absence of any true expression differences in the original simulated data. For the main experiments, we varied the prediction bias parameter *µ*. We used random seeds of 444 in all sampling steps.

### 5.7 Cell type annotation procedure for mouse hippocampus seqFISH dataset

We preprocessed the data using a standard Scanpy pipeline. Starting with the counts matrix, we normalized the data using pp.normalize_total() with default settings, log-transformed the data using pp.log1p(), and scaled the data with pp.scale(). We computed principal components and a neighbors graph using tl.pca() followed by pp.neighbors() with 20 principal components. Finally, we performed Leiden clustering using tl.leiden() with resolution of 0.3, which yielded 5 cell clusters. We used tl.rank_genes_groups() with the Wilcoxon method to identify the top five marker genes for each cell cluster and manually identified the clusters using those markers. In total, we identified endothelial cells, oligodendrocytes, astrocytes, and two neuron clusters.

### 5.8 Cell type annotation procedure for mouse VISP MERFISH dataset

We preprocessed the data using a standard Scanpy pipeline. Starting with the counts matrix, we normalized the data using pp.normalize_total() with default settings, log-transformed the data using pp.log1p(), and scaled the data with pp.scale(). We computed principal components and a neighbors graph using tl.pca() followed by pp.neighbors() with 20 principal components. Finally, we performed Leiden clustering using tl.leiden() with resolution of 0.1, which yielded 11 cell clusters. We used tl.rank_genes_groups() with the Wilcoxon method to identify the top five marker genes for each cell cluster and manually identified the clusters using those markers. In total, we identified endothelial cells, oligodendrocytes, astrocytes, and eight neuron-like cell clusters.

### 5.9 Anatomic region annotation procedure for Drosophila embryo dataset

We used the same preprocessing procedure as for the mouse VISP MERFISH dataset. We identified seven Leiden clusters and grouped them into four region labels based on their spatial localization with two “posterior” clusters, one “anterior” cluster, one “bottom” cluster, and three “middle” clusters.

### 5.10 Annotations for mouse somatosensory osmFISH dataset, mouse gastrulation seqFISH dataset, and axolotl telencephalon Stereo-seq dataset

We retrieved annotated class labels from publicly available metadata for these spatial transcriptomics datasets. For the mouse somatosensory osmFISH dataset, we retrieved both anatomic region (‘Region’) and cell type (‘ClusterName’) labels from the metadata available at http://linnarssonlab.org/osmFISH/osmFISH_SScortex_mouse_all_cells.loom. For the mouse gastrulation seqFISH dataset, we retrieved cell type (‘celltype_mapped_refined’) labels from the metadata available at https://content.cruk.cam.ac.uk/jmlab/SpatialMouseAtlas2020/ in metadata.Rds file for ‘embryo1’ and ‘z5’. For the axolotl telencephalon Stereo-seq dataset, we retrieved cell type (‘Annotation’) labels from the metadata available at https://db.cngb.org/stomics/artista/ for the Stage44.h5ad object file.

### 5.11 Replicate analysis for mouse gastrulation seqFISH dataset

To examine the reproducibility of TISSUE quantities across spatial transcriptomics replicates, we curated a replicate of the mouse gastrulation seqFISH dataset[37], that has not been previously included in benchmarking analyses for spatial gene expression prediction [33]. We mapped cell type (‘celltype_mapped_refined’) labels from the metadata available at https://content.cruk.cam.ac.uk/jmlab/SpatialMouseAtlas2020/ in metadata.Rds file for ‘embryo1’ and ‘z2’. For replication experiments, we utilized identical settings for TISSUE calibration, prediction interval calculation, and differential gene expression analysis as was used for the original dataset analysis.

### 5.12 Simulated prediction bias for clustering, visualization, and supervised learning

We generated synthetic data using SRTsim [72] for benchmarking TISSUE cell filtering and TISSUE WPCA approaches for improved performance on downstream analysis tasks. We used the reference-free SRTsim framework and generated a synthetic counts matrix using their native Shiny app. The data consists of two cell groups, referred to as ‘A’ and ‘B’, which were determined by manual drawing of a linear boundary between two spatial domains in the SRTsim Shiny application. There are 476 ‘A’ cells and 504 ‘B’ cells for a total of 980 cells. To generate counts, we followed the default SRTsim recommendations and used the zero-inflated negative binomial model and set the zero proportion to 0.05, dispersion to 0.5, and mean to 2. We simulated 1000 genes, consisting of 500 positive signal genes and 500 noise genes, where the positive signal genes had an average log fold change that was double in ‘B’ cells than in ‘A’ cells. We used a random seed of 444 for SRTsim.

To simulate prediction bias for cells in cell type ‘A’, we randomly sampled a proportion of cells in the ‘A’ group specified ‘mix-in’ proportion parameter and then for each gene and selected cell, we updated their expression level with a random uniform sample of ‘B’ cell expression levels for that gene. For the sampled cells, this simulated prediction bias shifts their predicted gene expression profiles to be more similar to that of cell type ‘B’ rather than cell type ‘A’. Finally, standard Gaussian noise was added to all other expression values for both cell types to simulated prediction noise. We used random seeds of 444 in all sampling steps.

### 5.13 Uncertainty-aware cell filtering for downstream tasks

Using the TISSUE prediction interval, we perform filtering of high-uncertainty cells to improve training/evaluation of supervised learning models, clustering, and data visualization. We approximate the prediction uncertainty using the width of the 67% prediction interval (equivalent to the asymmetric standard error). Then, we convert all uncertainty values to z-scores using the mean and standard deviation of expression for each gene in the data. For each cell, we assign a score equal to the average of its z-scores across all genes. The cells with the highest scores are removed from the filtered data. The threshold for removal is automatically determined using Otsu’s method, which finds a threshold that maximizes the variance between the filtered and unfiltered score sets. In the context of classification, we avoid inter-class differences in prediction uncertainty by performing this filtering procedure independently within each class.

### 5.14 Empirical evaluation of TISSUE cell filtering for downstream tasks

We used several evaluation metrics to quantify the improvement of TISSUE cell filtering over using the predicted gene expression (baseline) for a variety of common downstream analysis tasks. To ensure relatively balanced representation of classes, we used dataset and class label pairs that were restricted to the three classes with greatest prevalence. To generate the initial predicted spatial gene expression, we iteratively made predictions on held-out folds of genes using one of the specified prediction methods and with 10-fold cross-validation (see “Cross-validated spatial gene expression prediction” section for further details). For supervised learning (classification), we performed five-fold cross-validation where TISSUE cell filtering was applied independently on each train and test split. Within each fold, we fitted a logistic regression model on the train set using sklearn.linear model.LogisticRegression() with penalty=‘l1’ and solver=‘liblinear’. The model was evaluated on the test set and the classification accuracy, area under the receiver-operator curve, and macro F1 score are computed. These performance metrics were then averaged across the five folds. For clustering and visualization, we applied TISSUE cell filtering to the predicted gene expression data and then performed standardization and principal component analysis on the filtered data. For clustering, we then used k-means clustering with *k* = 3 on the top 15 principal components of the TISSUE-filtered data and measured clustering quality using the adjusted rand index with sklearn.metrics.adjusted_rand_score(). For visualization, we then fit a support vector classifier on the top 15 principal components of the TISSUE-filtered data using sklearn.svm.SVC() with kernel=‘linear’ and random state=444, and measured the accuracy of separation of classes, which we refer to as linear separability. For comparison, we repeated each of these procedures for the unfiltered/baseline predicted gene expression. These assessment procedures were applied independently for each spatial gene expression prediction method (Harmony, SpaGE, Tangram) within each dataset.

### 5.15 Dynamic visualization of the first two principal components with DynamicViz

We generated dynamic visualizations of the first two principal components for visual comparison of PCA on the measured spatial gene expression, PCA on the predicted spatial gene expression, and PCA on the TISSUE-filtered predicted spatial gene expression. We used DynamicViz (v.0.0.3) to center and rigidly align the cells across 20 two-dimensional PCA visualizations of the simulated datasets and visualized the resulting alignments using dynamicviz.viz.stacked(). Alignment was achieved on the subset of cells that overlapped between the reference and target visualizations for the TISSUE-filtered data. Robust visualizations can be consistently aligned across different replicates. We scored the variability of the resulting visualization by computing variance scores for each cell using dynamicviz.score.variance() with method=’global’.

### 5.16 Weighted PCA for uncertainty-aware tasks

As an alternative to TISSUE cell filtering, we implemented a weighted version of principal component analysis where each value in the gene expression matrix is assigned a scalar weight. We compute the weights according to the following steps. First, we compute the inverse of the TISSUE prediction interval width (i.e. 67% prediction interval upper bound minus lower bound). Then, we normalize these values for each gene by the mean value across that gene to correct for expression level differences between genes. Finally, we binarize these normalized values so that the top 80% of normalized values will have ten-fold higher relative weight than the bottom 20% of normalized values. These binary values are used as weights for Weighted Principal Component Analysis (WPCA). Alternatively, we have also implemented a weighting scheme where we simply take the log transform of the normalized inverse prediction interval widths, which provides comparable performance to the previously described weighting scheme (Fig. S7C). WPCA directly decomposes the weighted covariance matrix to obtain principal vectors, and then applies weighted least squares optimization to retrieve the principal components [18]. We used the implementation of WPCA in the wpca (v.0.1) Python package with default settings and weights set according to our specification. The TISSUE implementation of WPCA is customizable with user options for specifying different weighting parameters.

### 5.17 Sample processing for MERFISH dataset of mouse subventricular zone

A healthy 3-month old male C57BL/6 mouse was obtained from the National Institute on Aging (NIA) Aged Rodent colony. The mouse was habituated for at least two weeks at Stanford before use. It was housed at the ChEM-H/Neuroscience Vivarium at Stanford and their care was monitored by the Veterinary Service Center at Stanford University under the Institutional Animal Care and Use Committee protocol 8661.

The mouse was euthanized in a carbon dioxide chamber in the morning. The whole brain was dissected and immediately embedded in ice cold Tissue-Tek O.C.T. compound in a cryomold and placed on dry ice. Once the sample was frozen it was transferred and stored at −80^°^ C. The sample was shipped to Vizgen, Inc. in dry ice for processing. At Vizgen, the brain was sectioned to obtain coronal sections of the subventricular zone followed by MERFISH lab service, transcript count detection, and cell segmentation and allocation of counts to individual cells. The MERFISH dataset includes two consecutive coronal sections.

We curated a 140-gene panel for the MERFISH experiment. The panel included 2-5 known transcriptomic cell type markers for each of aNSC/NPCs, qNSC/astrocytes, neuroblasts, microglia, endothelial cells, oligodendrocytes, T cells, mural cells, ependymal cells, neurons, macrophages, and reactive astrocytes [12, 21]. Markers for aNSC/NPCs were *Hmgb2, Hmgn2, Ccnd2, Sox2*; markers for qNSC/astrocytes were *Aldoc, Clu, Mt3, Gfap, Id4*; markers for neuroblasts were *Tubb2b, Sox4, Tubb3, Dcx*. The remaining genes in the panel were related to neurogenesis, T cell activity, glycolysis, lipid metabolism, and aging.

### 5.18 Data preprocessing and labeling for MERFISH dataset of mouse subventricular zone

The MERFISH dataset was cropped to around the left and right lateral ventricles using rectangular bounding boxes. The raw counts were normalized by the volume of the cell segmentation. To remove doublets and segmentation artifacts, we filtered out the top 2% and bottom 2% of cells by total normalized expression. For initial clustering and visualization of the data, we further normalized total expression of all cells to 250 (scanpy.pp.normalize|_total() with target_sum=250), log-transformed with an added pseudocount (scanpy.pp.log1p()), and scaled to z-scores (scanpy.pp.scale() with max_value=10). We performed PCA using scanpy.tl.pca(), built a neighbors graph with scanpy.pp.neighbors(), obtained UMAP visualization with scanpy.tl.umap(), and performed Leiden clustering with scanpy.tl.leiden() with resolution=0.5. Using visualizations of the cell type markers in the MERFISH gene panel along with differential expression analysis, we manually identified nine cell types clusters including two neuron clusters, astrocytes, oligodendrocytes, endothelial cells, ependymal cells, microglia, oligodendrocyte progenitor cells, and an ambiguous cell type cluster that localized to the lateral ventricles. We further sub-clustered the ambiguous cell cluster using the same Leiden clustering restricted to cells in this cluster and recovered three sub-clusters. For spatial region labels, we manually selected vertical coordinate cutoffs that corresponded to dorsal and ventral regions outlined in previous studies [14].

### 5.19 Ambiguous sub-cluster identification for MERFISH dataset of mouse subventricular zone

To assist in the identification of the ambiguous cell sub-clusters, we used TISSUE to obtain uncertainty-aware SpaGE spatial gene expression predictions for additional cell type marker genes that were not in the 140-gene MERFISH panel. These included general NSC and qNSC/astrocyte markers (*Slc1a3, Nr2e1, Sox9, Vcam1, Hes5, Prom1, Thbs4*), aNSC/NPC markers (*Pclaf, H2afx, Rrm2, Insm1, Egfr, Prom1, Mcm2, Cdk1*), and neuroblast markers (*Stmn2, Dlx6os1, Igfbpl1, Sox11, Dlx1*). For prediction, we used a single-cell RNAseq dataset of the micro-dissected mouse subventricular zone [12]. To perform differential expression analysis, we used the TISSUE multiple imputation framework to perform two-sample t-tests to compare the expression of each predicted gene in one of the ambiguous sub-clusters in comparison to all other ambiguous sub-clusters. Final cell type identifications were made by considering the markers with statistically significant over-expression within each of the sub-clusters after Bonferroni multiple hypothesis correction.

### 5.20 Spatial region classifiers for MERFISH dataset of mouse subventricular zone

To train the regional SVZ classifiers, we performed TISSUE uncertainty-aware SpaGE spatial gene expression prediction of the whole-transcriptome (i.e. all genes in the paired single-cell RNAseq dataset) and obtained p-values for differential expression in dorsal versus ventral regions for each cell type (qNSC/astrocyte, aNSC/NPC, neuroblast) using the TISSUE multiple imputation t-test. For each cell type, we selected the top 20 most differentially expressed genes with the lowest TISSUE multiple imputation t-test p-values across the dorsal and ventral regions. Then, we trained cell type-specific penalized logistic regression models (sklearn.linear model.LogisticRegression() with penalty=‘l1’ and solver=‘liblinear’) to predict the regional origin of the cell from these 20 predicted gene expression features. The inputs were standardized before fitting the logistic regression model. We obtained class probabilities for each cell using 10-fold cross validation, training and evaluating an independent model on each train and test split. For comparison, we used either the TISSUE-filtered input with the 67% prediction interval width or unfiltered input in fitting and evaluating the classifiers. For each cell type and approach (TISSUE-filtered or baseline unfiltered predicted expression), we measured the performance of the classifiers using the F1 score, accuracy, area under the receiver-operator curve, and average precision score using the corresponding Scikit-learn implementations of these metrics.

## Data Availability

Intermediate data files can be provided upon reasonable request. Raw data were accessed from existing benchmark datasets [33] and are also available from the following studies:

Mouse Hippocampus: Spatial transcriptomics (seqFISH) at https://content.cruk.cam.ac.uk/jmlab/SpatialMouseAtlas2020/; RNAseq (10X Chromium) at GSE158450 in GEO for ‘HIPP sc Rep1 10X sample’.

Mouse VISP: Spatial transcriptomics (MERFISH) at https://github.com/spacetx-spacejam/data/; RNAseq (Smart-seq) at https://portal.brain-map.org/atlases-and-data/rnaseq/mouse-v1-and-alm-smart-seq for mouse primary visual cortex (VISp).

Mouse Prefrontal Cortex (PC): Spatial transcriptomics (STARmap) at ‘20180419 BZ9 control’ in https://www.starmapresources.com/data; RNAseq (10X Chromium) at GSE158450 in GEO for ‘PFC sc Rep2 10X’.

Human Middle Temporal Gyrus (MTG): Spatial transcriptomics (ISS) at https://github.com/spacetx-spacejam/data; RNAseq (Smart-seq) at https://portal.brain-map.org/atlases-and-data/rnaseq/human-mtg-smart-seq.

Mouse VISP: Spatial transcriptomics (ISS) at https://github.com/spacetx-spacejam/data; RNAseq (Smart-seq) at https://portal.brain-map.org/atlases-and-data/rnaseq/mouse-v1-and-alm-smart-seq for mouse primary visual cortex (VISp).

Drosophila Embryo: Spatial transcriptomics (FISH) at https://github.com/rajewsky-lab/distmap; RNAseq (Drop-seq) at GSE95025 in GEO.

Mouse Somatosensory Cortex (SC): Spatial transcriptomics (osmFISH) at http://linnarssonlab.org/osmFISH/ for cortical region subset; RNAseq (Smart-seq) at https://portal.brain-map.org/atlases-and-data/rnaseq/mouse-whole-cortex-and-hippocampus-smart-seq for mouse somatosensory cortex (SSp).

Mouse VISP: Spatial transcriptomics (ExSeq) at https://github.com/spacetx-spacejam/data; RNAseq (Smart-seq) at https://portal.brain-map.org/atlases-and-data/rnaseq/mouse-v1-and-alm-smart-seq for mouse primary visual cortex (VISp).

## Code Availability

The TISSUE Python package and associated code and documentation are available at https://github.com/sunericd/TISSUE, and all code for generating figures and analyses are separately available at https://github.com/sunericd/tissue-figures-and-analyses.

## Acknowledgements

Funding support was provided by Knight-Hennessy Scholars program (E.D.S.), Paul and Daisy Soros Fellowship for New Americans (E.D.S.), the National Science Foundation Graduate Research Fellowship Program (E.D.S.), Professor David Donoho at Stanford University (R.M.), NIH P01AG036695 (A.B.), NSF CAREER 1942926 (J.Z.), NIH P30AG059307 (J.Z.), 5RM1HG010023 (J.Z.) and grants from the Silicon Valley Foundation (J.Z.) and the Chan-Zuckerberg Initiative (J.Z.). We would like to thank L. Xu, O. Zhou, and M. Yuksekgonul for helpful discussions.

## Author’s contributions

E.D.S. and J.Z. conceived of the study. E.D.S. designed and implemented the method and ran all associated analyses with J.Z. and R.M providing input. P.N.N. and A.B. provided samples for the mouse subventricular zone MERFISH dataset and input on associated analyses. E.D.S. prepared a draft of the manuscript. R.M., P.N.N., A.B., and J.Z. edited the manuscript.

## 6 Extended Data Figures

**Figure S1:**
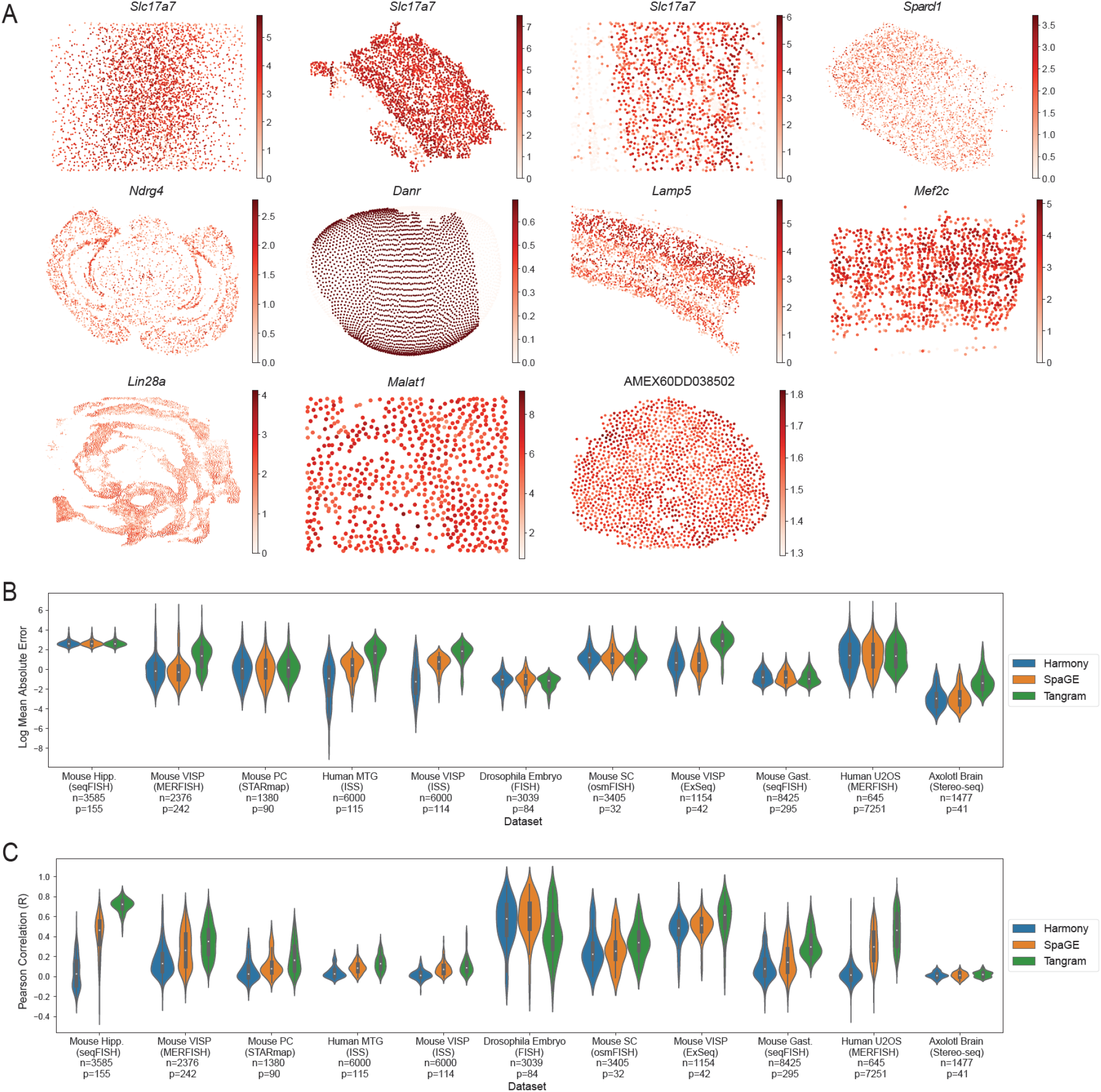
Overview of datasets and prediction performance. (A) Visualization of cells in the eleven spatial transcriptomics datasets colored by the expression of the highest-expressed gene in each respective dataset. (B-C) Performance of all three gene prediction methods (Harmony, SpaGE, Tangram) on all datasets as measured by (B) gene-wise mean absolute error between predicted and actual gene expression over 10-fold cross-validation, and (C) gene-wise Pearson correlation between predicted and actual gene expression over 10-fold cross-validation. Shown also are the number of cells (*n*) in the spatial transcriptomics datasets and the number of genes (*p*) shared between spatial and RNAseq datasets. In panels B-C, the inner box corresponds to quartiles of the metrics and the whiskers span up to 1.5 times the interquartile range of the metrics.

**Figure S2:**
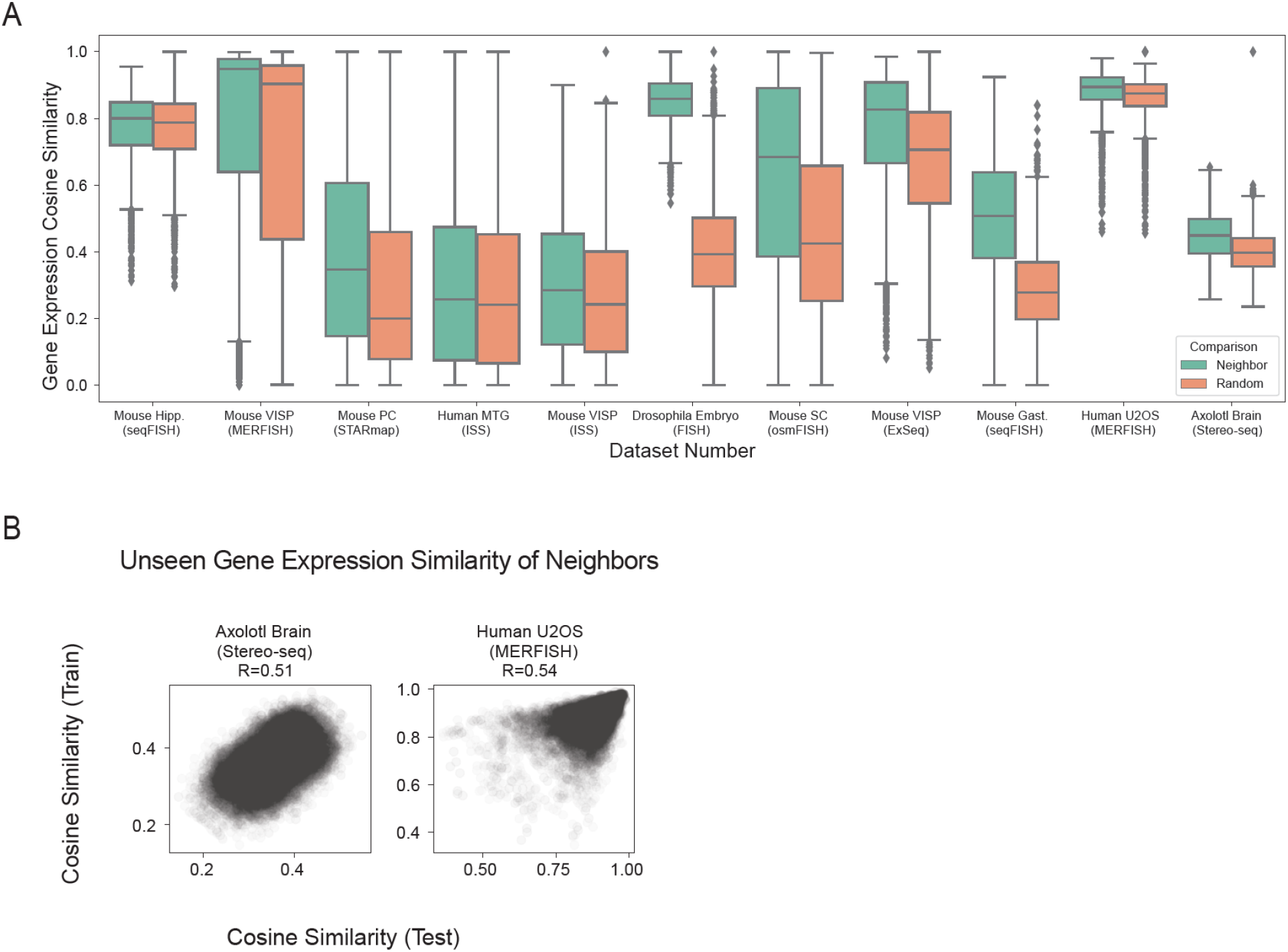
Evidence of gene expression similarity between spatial neighbors. (A) Cosine similarity of gene expression profiles for 250 cells paired with all their neighbors in the TISSUE spatial graph compared to pairings with randomly drawn cells across all eleven spatial transcriptomics datasets. The boxplot corresponds to the quartiles of the cosine similarity measurements. The center line corresponds to median cosine similarity, which was strictly higher in the neighbor-paired comparisons than the random-paired comparisons across all datasets. Whiskers span up to 1.5 times the interquartile range of the metrics and values outside this range are shown as dots. (B) Scatter plots of the cosine similarities of gene expression profiles for 250 cells paired with their neighbors for either the training gene set or the test gene set determined by random train-test split of all genes (50% train, 50% test). Shown are cosine similarity pairs for 10 train-test splits for the two datasets with the most genes.

**Figure S3:**
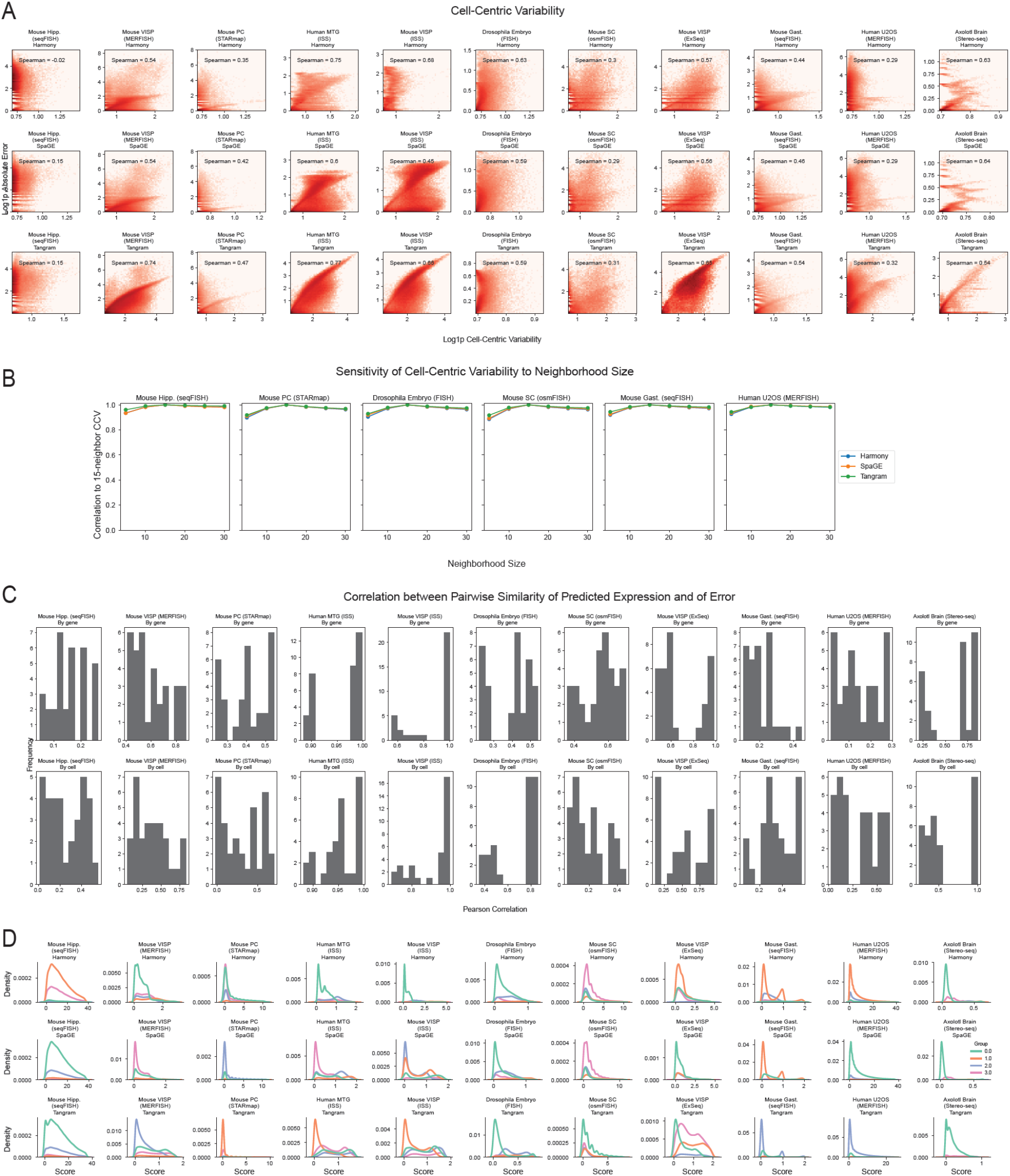
Cell-centric variability and calibration score distributions for individual datasets and prediction methods. (A) Correlation of cell-centric variability and absolute prediction error shown individually for each dataset and prediction method combination computed over 10-fold cross-validation. Log density with added pseudocount (Log1p) is shown by color, with a maximum of 1000 cells and 300 genes sampled from each dataset to provide more uniform representation. (B) Pearson correlation of all cell-centric variability measures obtained for different numbers of neighbors in building the TISSUE spatial graph compared to the default setting of 15 neighbors. Shown are results across six representative datasets. (C) Histograms showing the distribution of Pearson correlations between either gene-wise or cell-wise similarities of prediction errors and similarities of predicted expression values across all spatial transcriptomic datasets. (D) Distribution of TISSUE calibration scores shown individually for each dataset and prediction method combination ((*k*_*g*_, *k*_*c*_) = (4, 1)). Details on each dataset and prediction method can be found in Methods.

**Figure S4:**
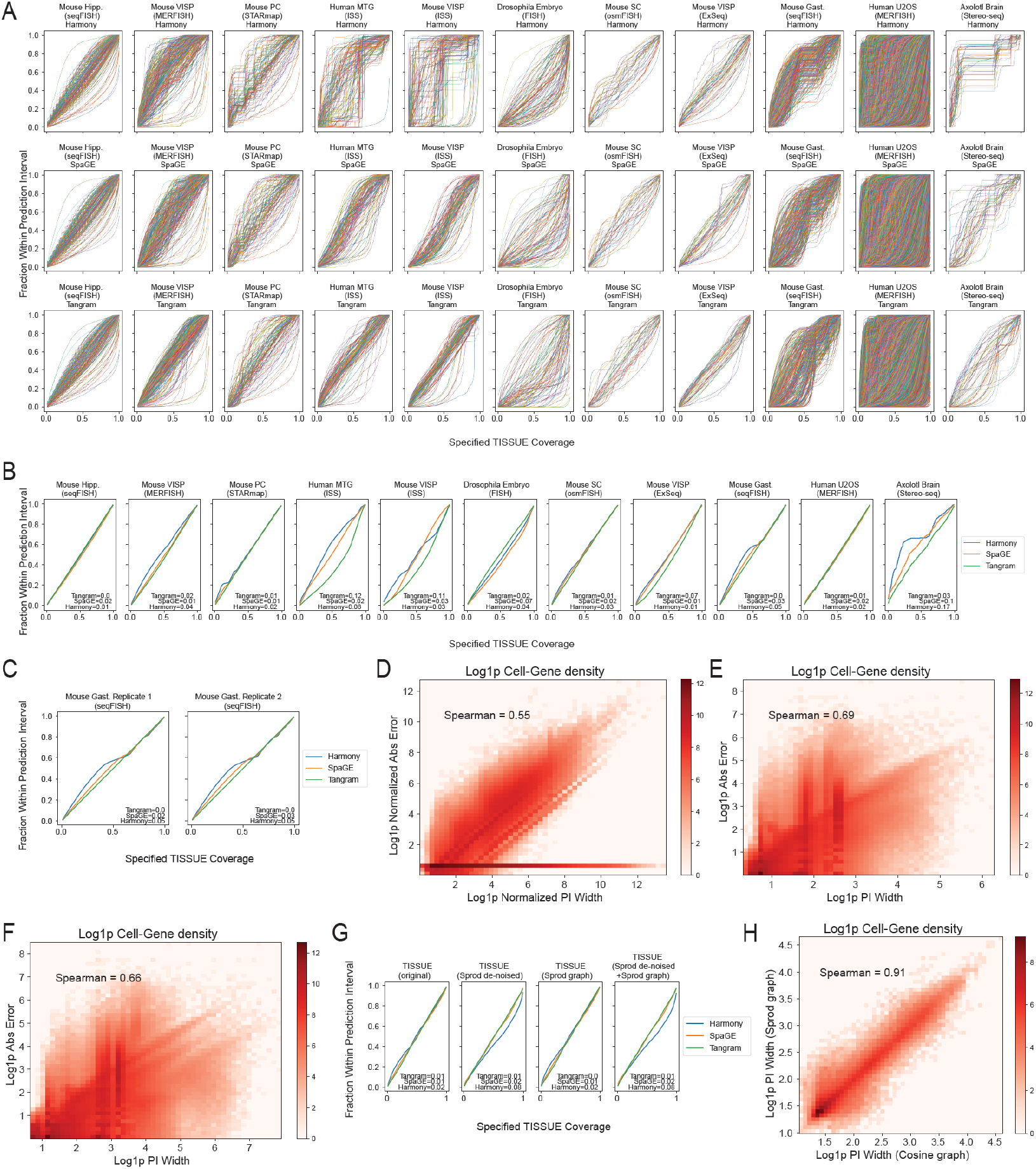
Further evaluation of TISSUE prediction intervals. (A) Gene-level calibration curves for TISSUE prediction intervals showing empirical coverage as a function of the specified confidence level across 10-fold cross-validation. Each line corresponds to an independent gene in the spatial transcriptomics dataset. (B-C) Calibration curves for TISSUE prediction intervals showing empirical coverage as a function of the specified confidence level across 10-fold cross-validation (B) under automated setting of (*k*_*g*_, *k*_*c*_) for stratified grouping; and (C) for two technical replicates of the mouse gastrulation seqFISH dataset with (*k*_*g*_, *k*_*c*_) = (4, 1). The calibration error is annotated for each prediction method (see Methods). (D-F) Correlation plots across all dataset and prediction method combinations computed over 10-fold cross-validation for (D) the 67% prediction interval width and absolute prediction error, both normalized by the absolute value of the predicted expression; (E) 50% prediction interval width and absolute prediction error; (F) 80% prediction interval width and absolute prediction error. Log density with added pseudocount (Log1p) is shown by color, with a maximum of 1000 cells and 300 genes sampled from each dataset to provide more uniform representation. (G) Calibration curves for TISSUE prediction intervals showing empirical coverage as a function of the specified confidence level across 10-fold cross-validation for the mouse somatosensory cortex osmFISH dataset with different combinations of Sprod de-noising or Sprod-based spatial similarity graph instead of the TISSUE spatial neighbors graph. The calibration error is annotated for each prediction method (see Methods). (H) Correlation plot of 67% prediction interval width with TISSUE spatial neighbors graph with cosine similarity weighting and 67% prediction interval width with Sprod similarity graph and weighting for the mouse somatosensory cortex osmFISH dataset and all prediction methods computed over 10-fold cross-validation.

**Figure S5:**
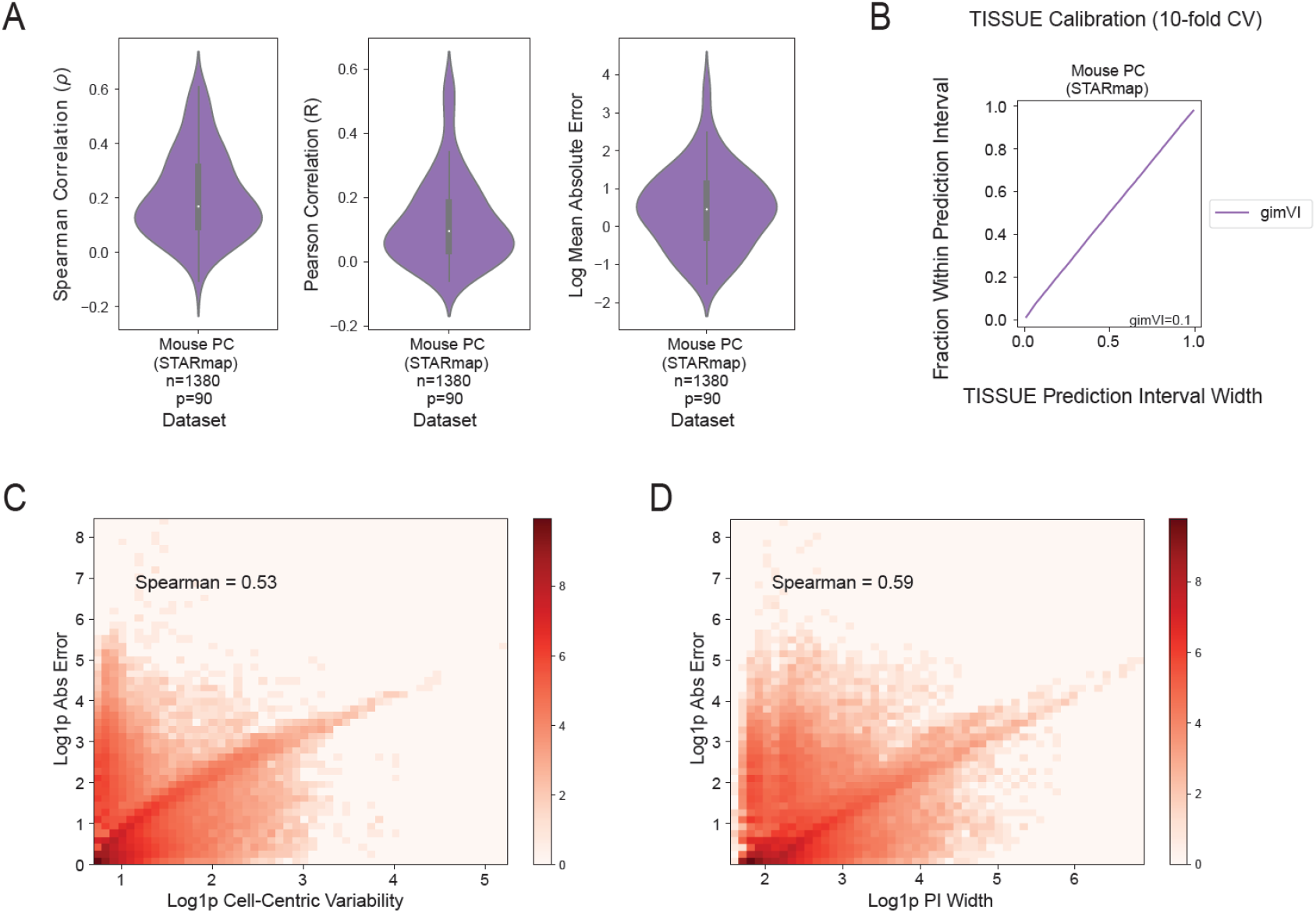
TISSUE results using gimVI for spatial gene expression prediction. (A) Violin plots showing the gene-wise performance of gimVI on predicting the gene expression of the mouse prefrontal cortex STARmap dataset using Spearman correlation (left), Pearson correlation (middle), and mean absolute error (right). The inner box corresponds to the quartiles of the bounds. (B) Calibration curve for TISSUE prediction intervals of gimVI showing empirical coverage as a function of the specified confidence level across 10-fold cross-validation for the mouse prefrontal cortex STARmap dataset with (*k*_*g*_, *k*_*c*_) = (4, 1). The calibration error is annotated for each prediction method (see Methods). (C) Correlation plot of cell-centric variability and the mean absolute prediction error of gimVI on the mouse prefrontal cortex STARmap dataset. (D) Correlation plot of 67% prediction interval width and the mean absolute prediction error of gimVI on the mouse prefrontal cortex STARmap dataset. Log density with added pseudocount (Log1p) is shown by color.

**Figure S6:**
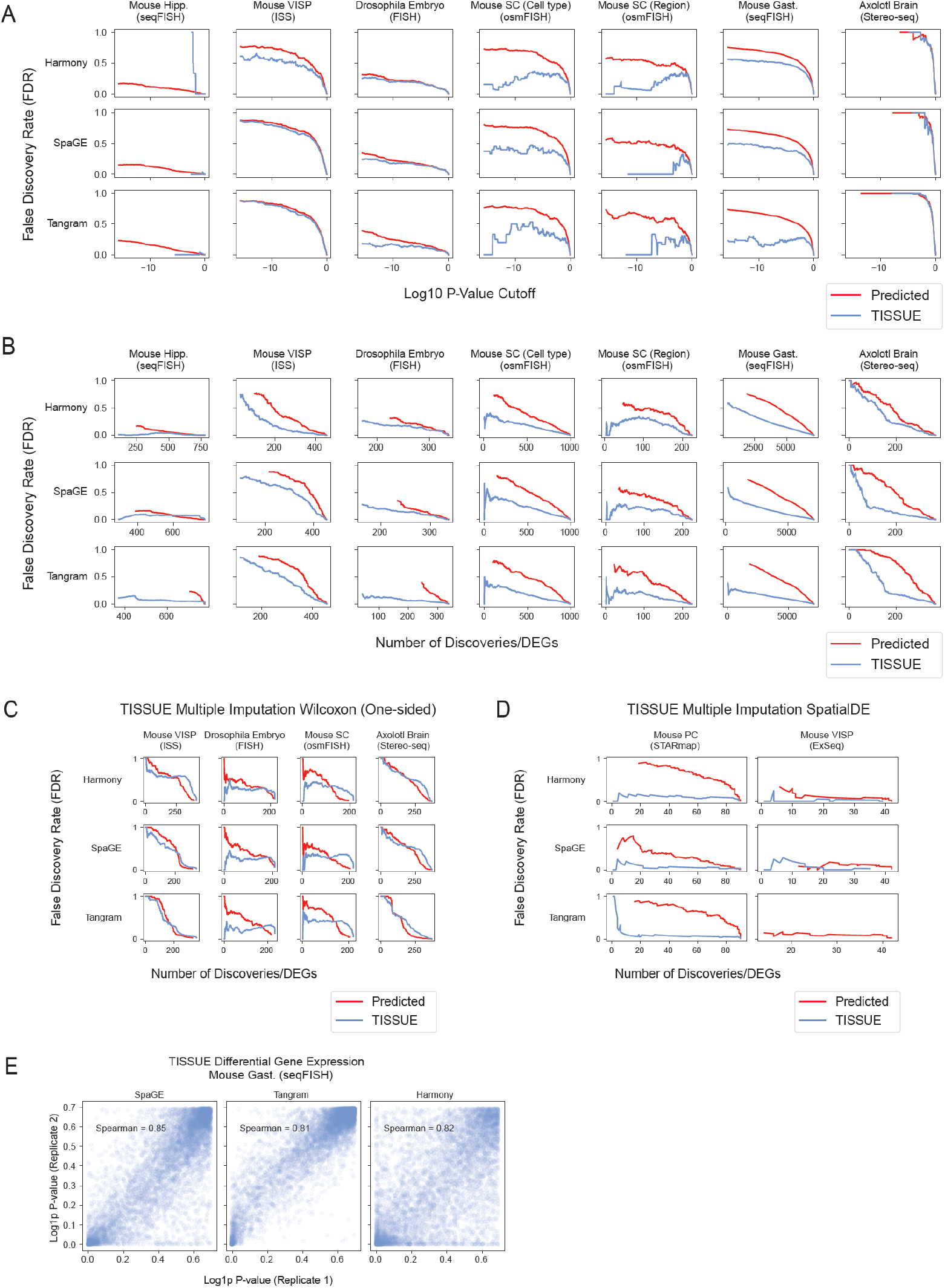
Additional differential gene expression analysis with TISSUE. (A) False discovery rate of differentially expressed genes between cell type or anatomic region labels (one versus all approach) as a function of the p-value threshold for significance and with (*k*_*g*_, *k*_*c*_) = (4, 1) settings for stratified grouping.(B) False discovery rate of differentially expressed genes between cell type or anatomic region labels (one versus all approach) as a function of the number of discoveries and with automated stratified grouping. (C) False discovery rate of differentially expressed genes between cell type or anatomic region labels (one versus all approach) as a function of the number of discoveries and with (*k*_*g*_, *k*_*c*_) = (4, 1) settings for stratified grouping for the alternative TISSUE multiple imputation framework using the “greater than” one-sided, two-sample Wilcoxon/Mann-Whitney test. (D) False discovery rate of spatially variable genes as a function of the number of discoveries and with (*k*_*g*_, *k*_*c*_) = (4, 1) settings for stratified grouping for the alternative TISSUE multiple imputation framework using the SpatialDE test. (E) Correlation plot of the log p-values obtained from the TISSUE multiple imputation t-test framework between two technical replicates of the mouse gastrulation seqFISH dataset.

**Figure S7:**
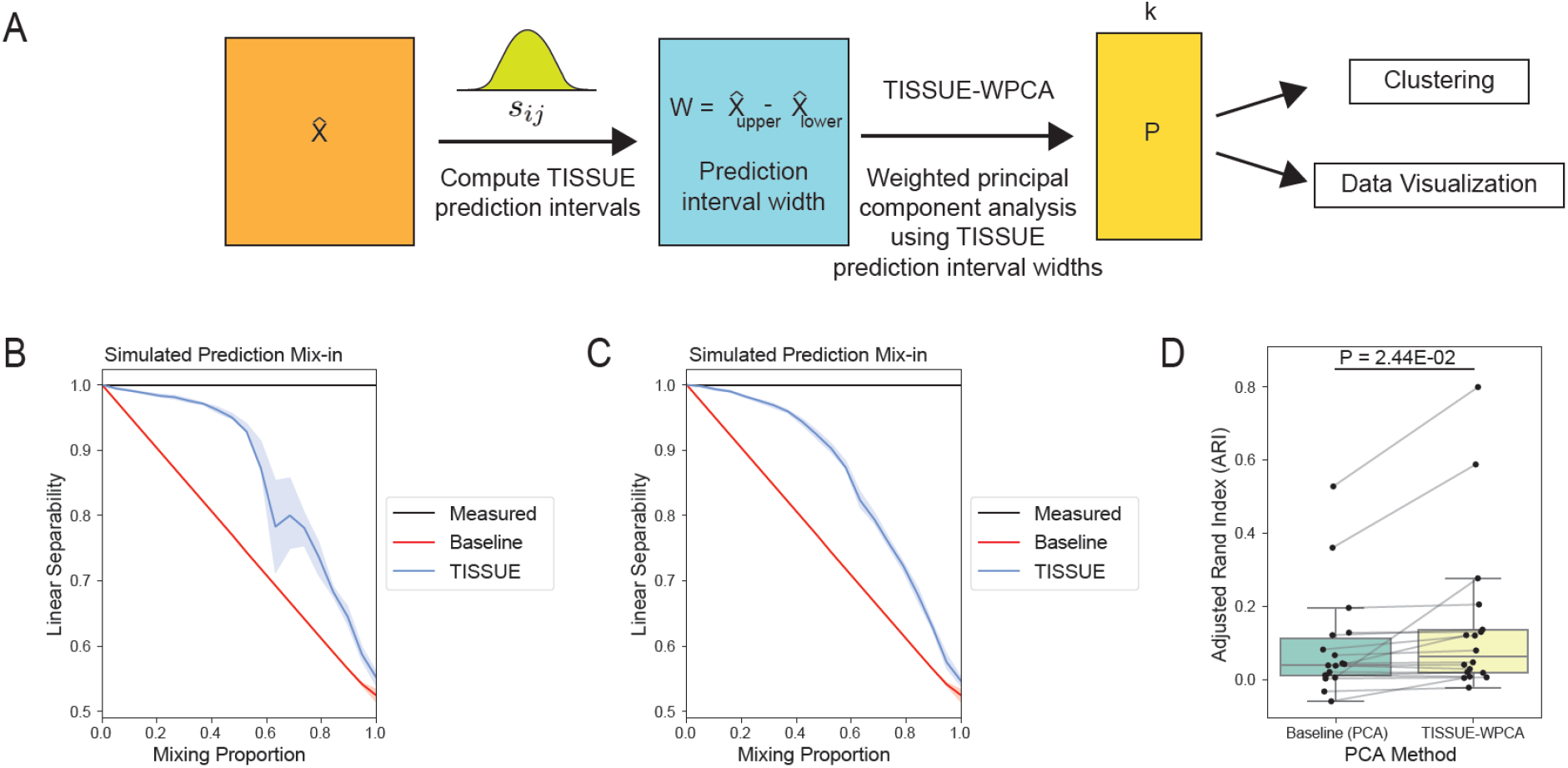
Uncertainty-aware clustering and label separation with TISSUE-WPCA. (A) Schematic illustration of the weighted principal component analysis (WPCA) pipeline where the inverse TISSUE prediction interval width is used to obtain principal components from WPCA, which are then used for downstream tasks of clustering and label separation. (B) Linear separability measured as the binary classification accuracy of a linear kernel support vector classifier fitted on the two cell clusters in the simulated spatial transcriptomics data as a function of the simulated mix-in proportion. The classifier was trained on the top 15 principal components obtained from the measured gene expression profiles with PCA, predicted gene expression profiles with PCA, and predicted gene expression profiles with TISSUE-WPCA. For TISSUE-WPCA, weights were determined by binarizing the inverse normalized 67% prediction interval width (see Methods). Results were obtained using automated stratified grouping. Bands represent the interquartile range and solid line denotes the median linear separability across 20 simulated datasets. (C) Same as in panel B except with TISSUE-WPCA weighting using the log-transformed inverse normalized 67% prediction interval width. (D) Adjusted rand index (ARI) for k-means clustering (*k* = 3) on the top 15 principal components obtained from PCA on the predicted expression or TISSUE-WPCA on the predicted gene expression for six real spatial transcriptomics dataset and labe pairings and all prediction methods. P-value was computed using a paired two-sample t-test. The box corresponds to quartiles of the metrics and the whiskers span up to 1.5 times the interquartile range of the metrics.

**Figure S8:**
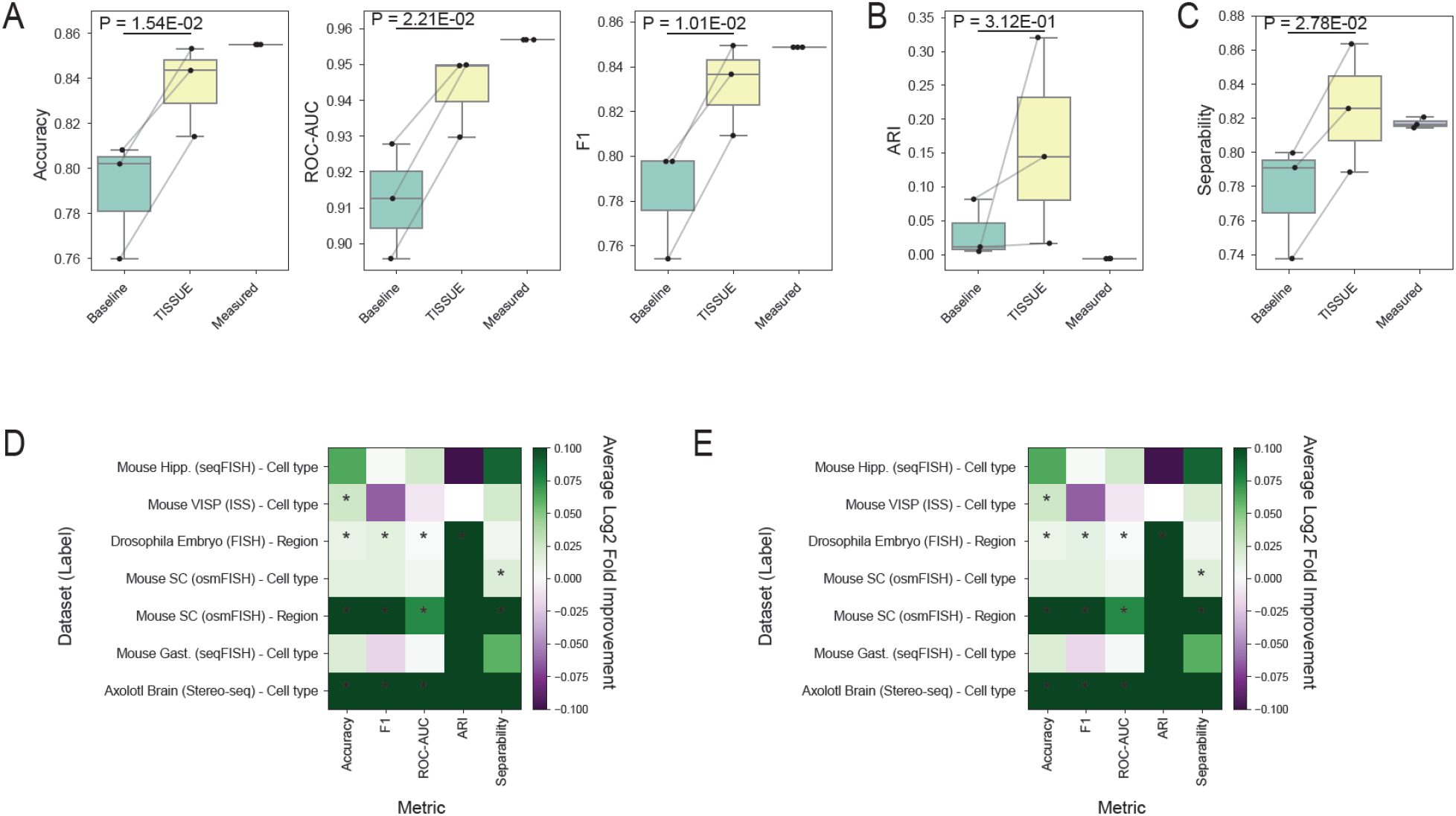
Additional experiments for uncertainty-aware supervised learning, clustering, and visualization. (A-C) Downstream task performance metrics for the three most prominent anatomic region class labels for the mouse somatosensory osmFISH dataset. Shown are metrics for all three prediction methods with automated stratified grouping settings. P-value was computed using a paired two-sample t-test. The box corresponds to quartiles of the metrics and the whiskers span up to 1.5 times the interquartile range of the metrics. (A) Accuracy, F1 score, and ROC-AUC (receiver-operator characteristic area under the curve) metrics for logistic regression models trained on the predicted gene expression, TISSUE-filtered predicted gene expression, or measured gene expression for classification. (B) Adjusted rand index (ARI) for k-means clustering (*k* = 3) on the top 15 principal components obtained from the predicted gene expression, TISSUE-filtered predicted gene expression, or measured gene expression for classification. (C) Linear separability measured as classification accuracy of linear kernel support vector classifier fitted on the top 15 principal components obtained from the predicted gene expression, TISSUE-filtered predicted gene expression, or measured gene expression for classification. (D) Log2 fold change in improvement of performance metrics using TISSUE-filtered PCA in lieu of regular PCA on predicted expression for supervised learning (logistic regression models trained to predict class labels; Accuracy, F1, ROC-AUC), clustering (k-means clustering on top 15 principal components; Adjusted Rand Index (ARI)), and visualization (linear separability measured as the classification accuracy of a linear kernel support vector classifier fitted on the top 15 principal components) for the top three classes across all dataset and class label combinations. Results were obtained using the 50% prediction interval width for filtering. (E) Same as panel E except with the 80% prediction interval width for filtering.

**Figure S9:**
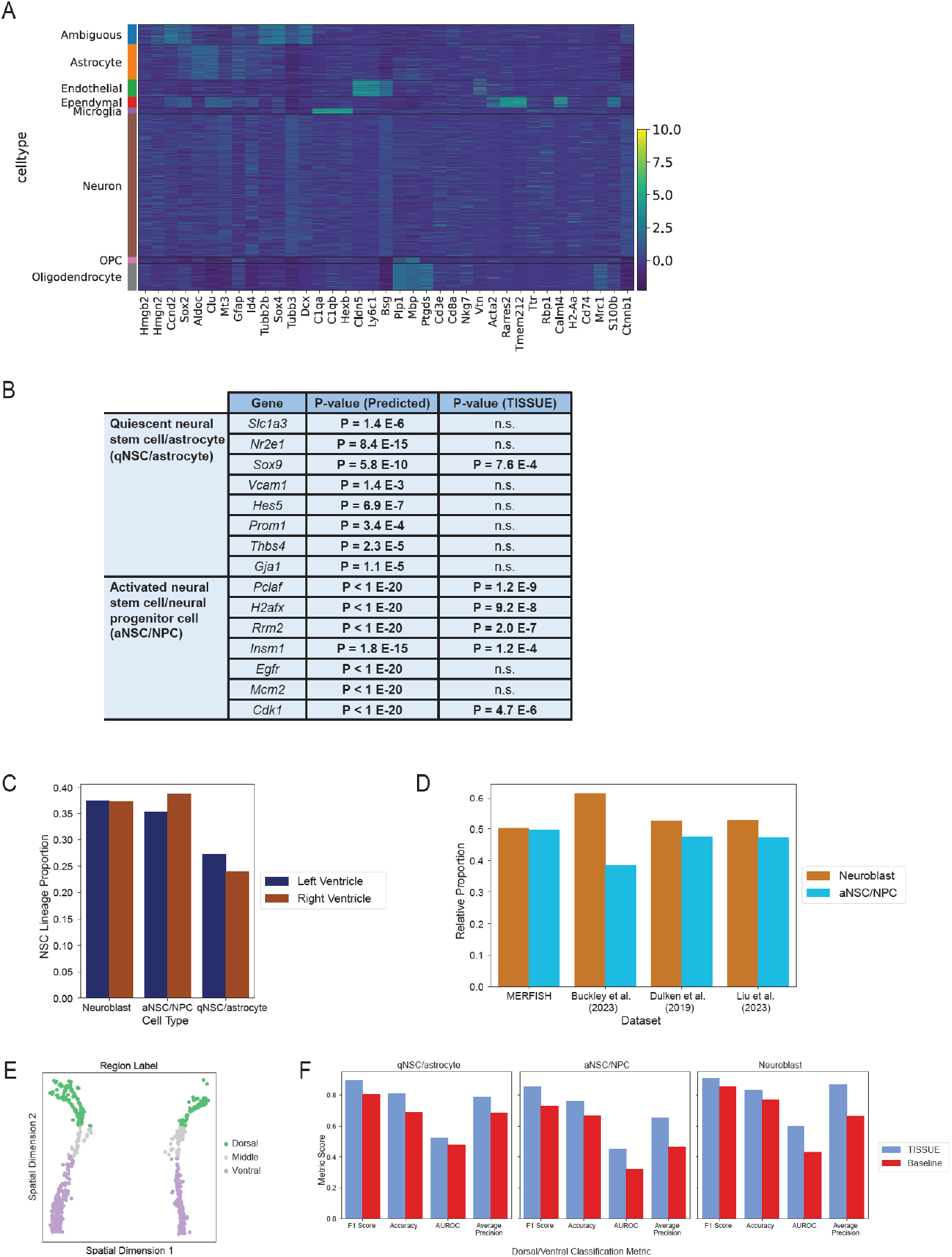
TISSUE is necessary to identify ambiguous NSC lineage subtype. (A) Heatmap of the scaled log-normalized gene expression of original cell type markers in the adult mouse subventricular zone MER-FISH dataset for each of the identified cell type clusters. The Ambiguous cell type cluster in the first row exhibits high expression of qNSC/astrocyte, aNSC/NPC, and neuroblast markers. (B) Additional predicted marker genes for the second ambiguous subcluster are differentially expressed for all qNSC/astrocyte and aNSC/NPC markers under traditional hypothesis testing with two-sample t-test on the predicted gene expression (Predicted). With TISSUE multiple imputation t-test, there are substantially more aNSC/NPC markers that are differentially over-expressed in the ambiguous subcluster (TISSUE), permitting identification of this subcluster as an aNSC/NPC subtype cluster. P-values are shown for all predicted marker genes with significance threshold of Bonferroni-adjusted *p <* 0.1. (C) Relative proportion of each of the three TISSUE-identified subtypes in the neural stem cell lineage cluster for either the left or right lateral ventricle. (D) Relative proportions of aNSC/NPC and neuroblast populations across the MERFISH dataset and three single-cell RNAseq datasets of the mouse subventricular zone. The qNSC/astrocyte proportions were not compared since they were aggregated with astrocytes of the striatum in the single-cell RNAseq datasets. (E) Spatial visualization of the cells in the neural stem cell lineage cluster colored by dorsal or ventral spatial location labels. (F) Dorsal versus ventral classification performance of TISSUE-filtered penalized logistic regression models and baseline unfiltered penalized logistic regression models evaluated using 10-fold cross-validation across F1 score, accuracy, area under the receiver-operator curve, and average precision.

**Figure S10:**
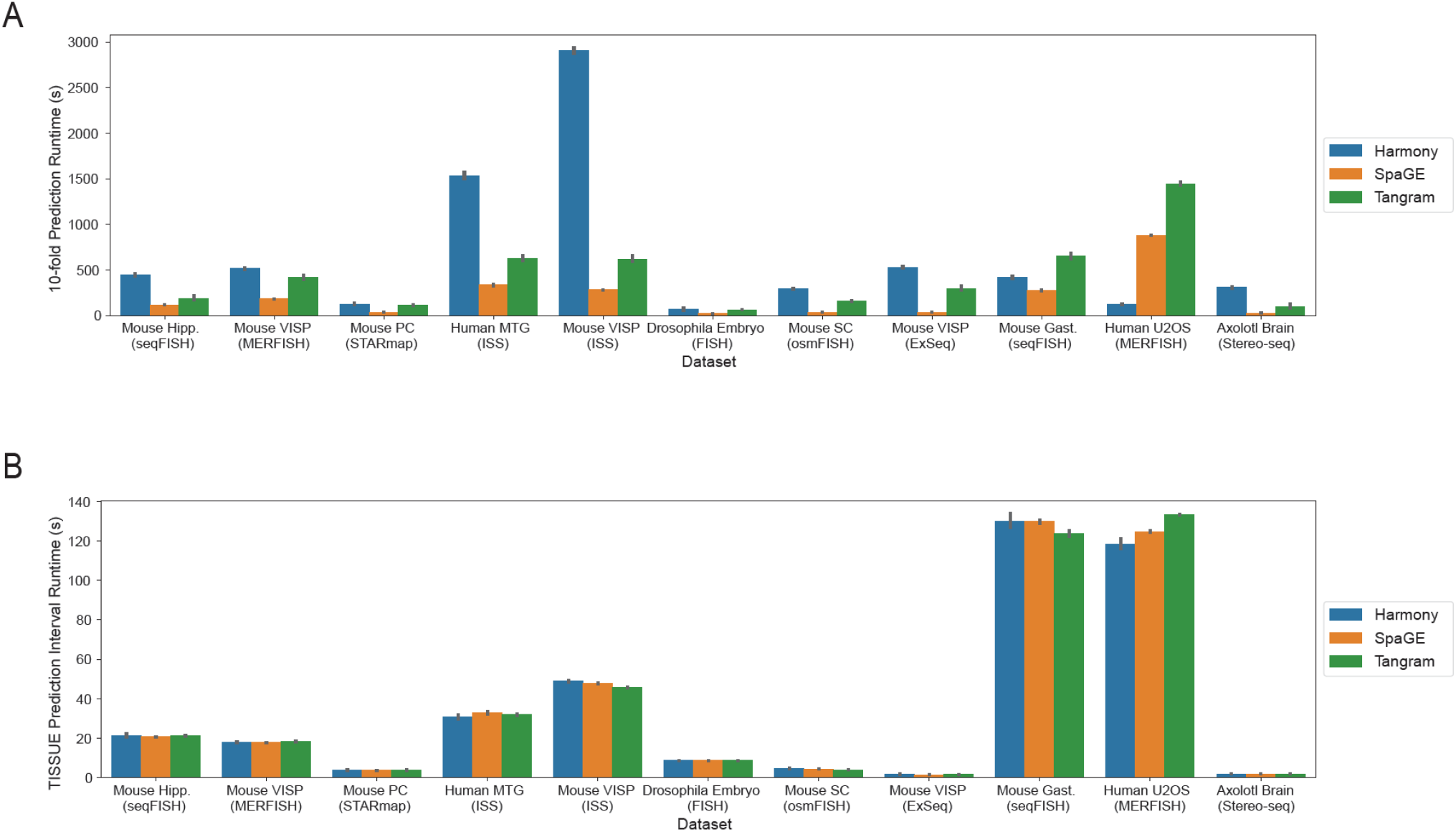
Computational runtime for TISSUE. (A) Bar plots of total runtimes for spatial gene expression prediction computations over 10 predictions to generate estimated predictions on all calibration genes. Error bars correspond to 95% confidence interval over an outer 10-fold cross-validation. (B) Bar plots of total runtimes for TISSUE prediction interval calculation including computation of cell-centric variability and calibration score sets. Error bars correspond to 95% confidence interval over an outer 10-fold cross-validation.

## References

[1] Tamim Abdelaal, Soufiane Mourragui, Ahmed Mahfouz, and Marcel J T Reinders. SpaGE: Spatial Gene Enhancement using scRNA-seq. Nucleic Acids Research, 48(18):e107, October 2020.

[2] William E. Allen, Timothy R. Blosser, Zuri A. Sullivan, Catherine Dulac, and Xiaowei Zhuang. Molecular and spatial signatures of mouse brain aging at single-cell resolution. Cell, 186(1):194–208.e18, January 2023. Publisher: Elsevier.

[3] Paul D. Allison. Missing Data.SAGE Publications, Inc., 2002.

[4] Shahar Alon, Daniel R. Goodwin, Anubhav Sinha, Asmamaw T. Wassie, Fei Chen, Evan R. Daugharthy, Yosuke Bando, Atsushi Kajita, Andrew G. Xue, Karl Marrett, Robert Prior, Yi Cui, Andrew C. Payne, Chun-Chen Yao, Ho-Jun Suk, Ru Wang, Chih-Chieh (Jay) Yu, Paul Tillberg, Paul Reginato, Nikita Pak, Songlei Liu, Sukanya Punthambaker, Eswar P. R. Iyer, Richie E. Kohman, Jeremy A. Miller, Ed S. Lein, Ana Lako, Nicole Cullen, Scott Rodig, Karla Helvie, Daniel L. Abravanel, Nikhil Wagle, Bruce E. Johnson, Johanna Klughammer, Michal Slyper, Julia Waldman, Judit Jané-Valbuena, Orit Rozenblatt-Rosen, Aviv Regev, IMAXT Consortium19, George M. Church, Adam H. Marblestone, and Edward S. Boyden. Expansion sequencing: Spatially precise in situ transcriptomics in intact biological systems. Science, 371(6528):eaax2656, January 2021. Publisher: American Association for the Advancement of Science.

[5] Arturo Alvarez-Buylla and Jose Manuel Garciia-Verdugo. Neurogenesis in Adult Subventricular Zone. Journal of Neuroscience, 22(3):629–634, February 2002. Publisher: Society for Neuroscience Section: MINI-REVIEW.

[6] Jonathan Alvarsson, Staffan Arvidsson McShane, Ulf Norinder, and Ola Spjuth. Predicting With Confidence: Using Conformal Prediction in Drug Discovery. Journal of Pharmaceutical Sciences, 110(1):42–49, January 2021.

[7] Anastasios N. Angelopoulos and Stephen Bates. A Gentle Introduction to Conformal Prediction and Distribution-Free Uncertainty Quantification, September 2022. arXiv:2107.07511 [cs, math, stat].

[8] Benedetta Artegiani, Anna Lyubimova, Mauro Muraro, Johan H. van Es, Alexander van Oudenaarden, and Hans Clevers. A Single-Cell RNA Sequencing Study Reveals Cellular and Molecular Dynamics of the Hippocampal Neurogenic Niche. Cell Reports, 21(11):3271–3284, December 2017.

[9] Michaela Asp, Stefania Giacomello, Ludvig Larsson, Chenglin Wu, Daniel Fürth, Xiaoyan Qian, Eva Wärdell, Joaquin Custodio, Johan Reimegård, Fredrik Salmén, Cecilia Österholm, Patrik L. Ståhl, Erik Sundström, Elisabet Åkesson, Olaf Bergmann, Magda Bienko, Agneta Månsson-Broberg, Mats Nilsson, Christer Sylvén, and Joakim Lundeberg. A Spatiotemporal Organ-Wide Gene Expression and Cell Atlas of the Developing Human Heart. Cell, 179(7):1647–1660.e19, December 2019.

[10] Tommaso Biancalani, Gabriele Scalia, Lorenzo Buffoni, Raghav Avasthi, Ziqing Lu, Aman Sanger, Neriman Tokcan, Charles R. Vanderburg, Åsa Segerstolpe, Meng Zhang, Inbal Avraham-Davidi, Sanja Vickovic, Mor Nitzan, Sai Ma, Ayshwarya Subramanian, Michal Lipinski, Jason Buenrostro, Nik Bear Brown, Duccio Fanelli, Xiaowei Zhuang, Evan Z. Macosko, and Aviv Regev. Deep learning and alignment of spatially resolved single-cell transcriptomes with Tangram. Nature Methods, 18(11):1352–1362, November 2021. Number: 11 Publisher: Nature Publishing Group.

[11] A. Sina Booeshaghi, Zizhen Yao, Cindy van Velthoven, Kimberly Smith, Bosiljka Tasic, Hongkui Zeng, and Lior Pachter. Isoform cell-type specificity in the mouse primary motor cortex. Nature, 598(7879):195–199, October 2021. Number: 7879 Publisher: Nature Publishing Group.

[12] Matthew T. Buckley, Eric D. Sun, Benson M. George, Ling Liu, Nicholas Schaum, Lucy Xu, Jaime M. Reyes, Margaret A. Goodell, Irving L. Weissman, Tony Wyss-Coray, Thomas A. Rando, and Anne Brunet. Cell-type-specific aging clocks to quantify aging and rejuvenation in neurogenic regions of the brain. Nature Aging, 3(1):121–137, January 2023. Number: 1 Publisher: Nature Publishing Group.

[13] Zixuan Cang and Qing Nie. Inferring spatial and signaling relationships between cells from single cell transcriptomic data. Nature Communications, 11(1):2084, April 2020. Number: 1 Publisher: Nature Publishing Group.

[14] Arantxa Cebrian-Silla, Marcos Assis Nascimento, Stephanie A Redmond, Benjamin Mansky, David Wu, Kirsten Obernier, Ricardo Romero Rodriguez, Susana Gonzalez-Granero, Jose Manuel Garcia-Verdugo, Daniel A Lim, and Arturo Álvarez Buylla. Single-cell analysis of the ventricular-subventricular zone reveals signatures of dorsal and ventral adult neurogenesis. eLife, 10:e67436, July 2021. Publisher: eLife Sciences Publications, Ltd.

[15] Zayna Chaker, Paolo Codega, and Fiona Doetsch. A mosaic world: puzzles revealed by adult neural stem cell heterogeneity. WIREs Developmental Biology, 5(6):640–658, 2016. eprint: https://onlinelibrary.wiley.com/doi/pdf/10.1002/wdev.248.

[16] Kok Hao Chen, Alistair N. Boettiger, Jeffrey R. Moffitt, Siyuan Wang, and Xiaowei Zhuang. Spatially resolved, highly multiplexed RNA profiling in single cells. Science, 348(6233):aaa6090, April 2015. Publisher: American Association for the Advancement of Science.

[17] Simone Codeluppi, Lars E. Borm, Amit Zeisel, Gioele La Manno, Josina A. van Lunteren, Camilla I. Svensson, and Sten Linnarsson. Spatial organization of the somatosensory cortex revealed by osmFISH. Nature Methods, 15(11):932–935, November 2018. Number: 11 Publisher: Nature Publishing Group.

[18] L. Delchambre. Weighted principal component analysis: a weighted covariance eigendecomposition approach. Monthly Notices of the Royal Astronomical Society, 446(4):3545–3555, February 2015.

[19] Fiona Doetsch. A niche for adult neural stem cells. Current Opinion in Genetics & Development, 13(5):543–550, October 2003.

[20] Kangning Dong and Shihua Zhang. Deciphering spatial domains from spatially resolved transcriptomics with an adaptive graph attention auto-encoder. Nature Communications, 13(1):1739, April 2022.

[21] Ben W. Dulken, Matthew T. Buckley, Paloma Navarro Negredo, Naresha Saligrama, Romain Cayrol, Dena S. Leeman, Benson M. George, Stéphane C. Boutet, Katja Hebestreit, John V. Pluvinage, Tony Wyss-Coray, Irving L. Weissman, Hannes Vogel, Mark M. Davis, and Anne Brunet. Single-cell analysis reveals T cell infiltration in old neurogenic niches. Nature, 571(7764):205–210, July 2019. Number: 7764 Publisher: Nature Publishing Group.

[22] Adam Gayoso, Zöe Steier, Romain Lopez, Jeffrey Regier, Kristopher L. Nazor, Aaron Streets, and Nir Yosef. Joint probabilistic modeling of single-cell multi-omic data with totalVI. Nature Methods, 18(3):272–282, March 2021. Number: 3 Publisher: Nature Publishing Group.

[23] Daniel Gyllborg, Christoffer Mattsson Langseth, Xiaoyan Qian, Eunkyoung Choi, Sergio Marco Salas, Markus M Hilscher, Ed S Lein, and Mats Nilsson. Hybridization-based in situ sequencing (HybISS) for spatially resolved transcriptomics in human and mouse brain tissue. Nucleic Acids Research, 48(19):e112, November 2020.

[24] Rebecca D. Hodge, Trygve E. Bakken, Jeremy A. Miller, Kimberly A. Smith, Eliza R. Barkan, Lucas T. Graybuck, Jennie L. Close, Brian Long, Nelson Johansen, Osnat Penn, Zizhen Yao, Jeroen Eggermont, Thomas Höllt, Boaz P. Levi, Soraya I. Shehata, Brian Aevermann, Allison Beller, Darren Bertagnolli, Krissy Brouner, Tamara Casper, Charles Cobbs, Rachel Dalley, Nick Dee, Song-Lin Ding, Richard G. Ellenbogen, Olivia Fong, Emma Garren, Jeff Goldy, Ryder P. Gwinn, Daniel Hirschstein, C. Dirk Keene, Mohamed Keshk, Andrew L. Ko, Kanan Lathia, Ahmed Mahfouz, Zoe Maltzer, Medea McGraw, Thuc Nghi Nguyen, Julie Nyhus, Jeffrey G. Ojemann, Aaron Oldre, Sheana Parry, Shannon Reynolds, Christine Rimorin, Nadiya V. Shapovalova, Saroja Somasundaram, Aaron Szafer, Elliot R. Thomsen, Michael Tieu, Gerald Quon, Richard H. Scheuermann, Rafael Yuste, Susan M. Sunkin, Boudewijn Lelieveldt, David Feng, Lydia Ng, Amy Bernard, Michael Hawrylycz, John W. Phillips, Bosiljka Tasic, Hongkui Zeng, Allan R. Jones, Christof Koch, and Ed S. Lein. Conserved cell types with divergent features in human versus mouse cortex. Nature, 573(7772):61–68, September 2019. Number: 7772 Publisher: Nature Publishing Group.

[25] Andrew L. Ji, Adam J. Rubin, Kim Thrane, Sizun Jiang, David L. Reynolds, Robin M. Meyers, Margaret G. Guo, Benson M. George, Annelie Mollbrink, Joseph Bergenstråhle, Ludvig Larsson, Yunhao Bai, Bokai Zhu, Aparna Bhaduri, Jordan M. Meyers, Xavier Rovira-Clavé, S. Tyler Hollmig, Sumaira Z. Aasi, Garry P. Nolan, Joakim Lundeberg, and Paul A. Khavari. Multimodal Analysis of Composition and Spatial Architecture in Human Squamous Cell Carcinoma. Cell, 182(2):497–514.e22, July 2020.

[26] Ying Jin, Zhimei Ren, and Emmanuel J. Candès. Sensitivity analysis of individual treatment effects: A robust conformal inference approach. Proceedings of the National Academy of Sciences, 120(6):e2214889120, February 2023. Publisher: Proceedings of the National Academy of Sciences.

[27] Anoushka Joglekar, Andrey Prjibelski, Ahmed Mahfouz, Paul Collier, Susan Lin, Anna Katharina Schlusche, Jordan Marrocco, Stephen R. Williams, Bettina Haase, Ashley Hayes, Jennifer G. Chew, Neil I. Weisenfeld, Man Ying Wong, Alexander N. Stein, Simon A. Hardwick, Toby Hunt, Qi Wang, Christoph Dieterich, Zachary Bent, Olivier Fedrigo, Steven A. Sloan, Davide Risso, Erich D. Jarvis, Paul Flicek, Wenjie Luo, Geoffrey S. Pitt, Adam Frankish, August B. Smit, M. Elizabeth Ross, and Hagen U. Tilgner. A spatially resolved brain region- and cell type-specific isoform atlas of the postnatal mouse brain. Nature Communications, 12(1):463, January 2021.

[28] Nikos Karaiskos, Philipp Wahle, Jonathan Alles, Anastasiya Boltengagen, Salah Ayoub, Claudia Kipar, Christine Kocks, Nikolaus Rajewsky, and Robert P. Zinzen. The Drosophila embryo at single-cell transcriptome resolution. Science, 358(6360):194–199, October 2017. Publisher: American Association for the Advancement of Science.

[29] Rongqin Ke, Marco Mignardi, Alexandra Pacureanu, Jessica Svedlund, Johan Botling, Carolina Wählby, and Mats Nilsson. In situ sequencing for RNA analysis in preserved tissue and cells. Nature Methods, 10(9):857–860, September 2013. Number: 9 Publisher: Nature Publishing Group.

[30] Ilya Korsunsky, Nghia Millard, Jean Fan, Kamil Slowikowski, Fan Zhang, Kevin Wei, Yuriy Baglaenko, Michael Brenner, Po-ru Loh, and Soumya Raychaudhuri. Fast, sensitive and accurate integration of single-cell data with Harmony. Nature Methods, 16(12):1289–1296, December 2019. Number: 12 Publisher: Nature Publishing Group.

[31] P R Langer-Safer, M Levine, and D C Ward. Immunological method for mapping genes on Drosophila polytene chromosomes. Proceedings of the National Academy of Sciences of the United States of America, 79(14):4381–4385, July 1982.

[32] Jing Lei, Max G’Sell, Alessandro Rinaldo, Ryan J. Tibshirani, and Larry Wasserman. Distribution-Free Predictive Inference for Regression. Journal of the American Statistical Association, 113(523):1094–1111, July 2018. Publisher: Taylor & Francis eprint: 10.1080/01621459.2017.1307116.

[33] Bin Li, Wen Zhang, Chuang Guo, Hao Xu, Longfei Li, Minghao Fang, Yinlei Hu, Xinye Zhang, Xinfeng Yao, Meifang Tang, Ke Liu, Xuetong Zhao, Jun Lin, Linzhao Cheng, Falai Chen, Tian Xue, and Kun Qu. Benchmarking spatial and single-cell transcriptomics integration methods for transcript distribution prediction and cell type deconvolution. Nature Methods, 19(6):662–670, June 2022. Number: 6 Publisher: Nature Publishing Group.

[34] Christine Licht. New methods for generating significance levels from multiply-imputed data. PhD thesis, Otto-Friedrich-Universität Bamberg, Fakultät Sozial-und Wirtschaftswissenschaften, 2010.

[35] Roderick J. A. Little and Donald B. Rubin. Bayes and Multiple Imputation. In Statistical Analysis with Missing Data, pages 200–220. John Wiley & Sons, Ltd, 2002. Section: 10_eprint: https://onlinelibrary.wiley.com/doi/pdf/10.1002/9781119013563.ch10.

[36] Ling Liu, Soochi Kim, Matthew T. Buckley, Jaime M. Reyes, Jengmin Kang, Lei Tian, Mingqiang Wang, Alexander Lieu, Michelle Mao, Cristina Rodriguez-Mateo, Heather D. Ishak, Mira Jeong, Joseph C. Wu, Margaret A. Goodell, Anne Brunet, and Thomas A. Rando. Exercise reprograms the inflammatory landscape of multiple stem cell compartments during mammalian aging. Cell Stem Cell, 30(5):689–705.e4, May 2023.

[37] T. Lohoff, S. Ghazanfar, A. Missarova, N. Koulena, N. Pierson, J. A. Griffiths, E. S. Bardot, C.- H. L. Eng, R. C. V. Tyser, R. Argelaguet, C. Guibentif, S. Srinivas, J. Briscoe, B. D. Simons, A.-K. Hadjantonakis, B. Göttgens, W. Reik, J. Nichols, L. Cai, and J. C. Marioni. Integration of spatial and single-cell transcriptomic data elucidates mouse organogenesis. Nature Biotechnology, 40(1):74–85, January 2022. Number: 1 Publisher: Nature Publishing Group.

[38] Brian Long, Jeremy Miller, and The SpaceTx Consortium. SpaceTx: A Roadmap for Benchmarking Spatial Transcriptomics Exploration of the Brain, January 2023. arXiv:2301.08436 [q-bio].

[39] Romain Lopez, Achille Nazaret, Maxime Langevin, Jules Samaran, Jeffrey Regier, Michael I. Jordan, and Nir Yosef. A joint model of unpaired data from scRNA-seq and spatial transcriptomics for imputing missing gene expression measurements, May 2019. arXiv:1905.02269 [cs, q-bio, stat].

[40] Mohammad Lotfollahi, F. Alexander Wolf, and Fabian J. Theis. scGen predicts single-cell perturbation responses. Nature Methods, 16(8):715–721, August 2019. Number: 8 Publisher: Nature Publishing Group.

[41] Eric Lubeck, Ahmet F. Coskun, Timur Zhiyentayev, Mubhij Ahmad, and Long Cai. Single-cell in situ RNA profiling by sequential hybridization. Nature Methods, 11(4):360–361, April 2014. Number: 4 Publisher: Nature Publishing Group.

[42] Katharina Lust, Ashley Maynard, Tomás Gomes, Jonas Simon Fleck, J. Gray Camp, Elly M. Tanaka, and Barbara Treutlein. Single-cell analyses of axolotl telencephalon organization, neurogenesis, and regeneration. Science, 377(6610):eabp9262, September 2022. Publisher: American Association for the Advancement of Science.

[43] Andrea Marshall, Douglas G Altman, Roger L Holder, and Patrick Royston. Combining estimates of interest in prognostic modelling studies after multiple imputation: current practice and guidelines. BMC Medical Research Methodology, 9:57, July 2009.

[44] Kristen R. Maynard, Leonardo Collado-Torres, Lukas M. Weber, Cedric Uytingco, Brianna K. Barry, Stephen R. Williams, Joseph L. Catallini, Matthew N. Tran, Zachary Besich, Madhavi Tippani, Jennifer Chew, Yifeng Yin, Joel E. Kleinman, Thomas M. Hyde, Nikhil Rao, Stephanie C. Hicks, Keri Martinowich, and Andrew E. Jaffe. Transcriptome-scale spatial gene expression in the human dorsolateral prefrontal cortex. Nature Neuroscience, 24(3):425–436, March 2021. Number: 3 Publisher: Nature Publishing Group.

[45] Reuben Moncada, Dalia Barkley, Florian Wagner, Marta Chiodin, Joseph C. Devlin, Maayan Baron, Cristina H. Hajdu, Diane M. Simeone, and Itai Yanai. Integrating microarray-based spatial transcriptomics and single-cell RNA-seq reveals tissue architecture in pancreatic ductal adenocarcinomas. Nature Biotechnology, 38(3):333–342, March 2020.

[46] Noa Moriel, Enes Senel, Nir Friedman, Nikolaus Rajewsky, Nikos Karaiskos, and Mor Nitzan. NovoSpaRc: flexible spatial reconstruction of single-cell gene expression with optimal transport. Nature Protocols, 16(9):4177–4200, September 2021. Number: 9 Publisher: Nature Publishing Group.

[47] Lambda Moses and Lior Pachter. Museum of spatial transcriptomics. Nature Methods, pages 1–13, March 2022. Publisher: Nature Publishing Group.

[48] Soufiane Mourragui, Marco Loog, Mark A van de Wiel, Marcel J T Reinders, and Lodewyk F A Wessels. PRECISE: a domain adaptation approach to transfer predictors of drug response from preclinical models to tumors. Bioinformatics, 35(14):i510–i519, July 2019.

[49] Paloma Navarro Negredo, Robin W. Yeo, and Anne Brunet. Aging and Rejuvenation of Neural Stem Cells and Their Niches. Cell Stem Cell, 27(2):202–223, August 2020.

[50] Mor Nitzan, Nikos Karaiskos, Nir Friedman, and Nikolaus Rajewsky. Gene expression cartography. Nature, 576(7785):132–137, December 2019. Number: 7785 Publisher: Nature Publishing Group.

[51] Cameron Palmer and Itsik Pe’er. Bias Characterization in Probabilistic Genotype Data and Improved Signal Detection with Multiple Imputation. PLOS Genetics, 12(6):e1006091, June 2016. Publisher: Public Library of Science.

[52] Antonio Scialdone, Kedar N. Natarajan, Luis R. Saraiva, Valentina Proserpio, Sarah A. Teichmann, Oliver Stegle, John C. Marioni, and Florian Buettner. Computational assignment of cell-cycle stage from single-cell transcriptome data. Methods, 85:54–61, September 2015.

[53] Glenn Shafer and Vladimir Vovk. A Tutorial on Conformal Prediction. Journal of Machine Learning Research, 9(12):371–421, 2008.

[54] Sheel Shah, Eric Lubeck, Wen Zhou, and Long Cai. In Situ Transcription Profiling of Single Cells Reveals Spatial Organization of Cells in the Mouse Hippocampus. Neuron, 92(2):342–357, October 2016.

[55] Chen Shengquan, Zhang Boheng, Chen Xiaoyang, Zhang Xuegong, and Jiang Rui. stPlus: a reference-based method for the accurate enhancement of spatial transcriptomics. Bioinformatics, 37(Supplement 1):i299–i307, July 2021.

[56] Tabrez J. Siddiqui, Parisa Karimi Tari, Steven A. Connor, Peng Zhang, Frederick A. Dobie, Kevin She, Hiroshi Kawabe, Yu Tian Wang, Nils Brose, and Ann Marie Craig. An LRRTM4-HSPG Complex Mediates Excitatory Synapse Development on Dentate Gyrus Granule Cells. Neuron, 79(4):680–695, August 2013.

[57] Tim Stuart, Andrew Butler, Paul Hoffman, Christoph Hafemeister, Efthymia Papalexi, William M. Mauck, Yuhan Hao, Marlon Stoeckius, Peter Smibert, and Rahul Satija. Comprehensive Integration of Single-Cell Data. Cell, 177(7):1888–1902.e21, June 2019.

[58] Eric D. Sun, Rong Ma, and James Zou. Dynamic visualization of high-dimensional data. Nature Computational Science, 3(1):86–100, January 2023. Number: 1 Publisher: Nature Publishing Group.

[59] Valentine Svensson, Sarah A. Teichmann, and Oliver Stegle. SpatialDE: identification of spatially variable genes. Nature Methods, 15(5):343–346, May 2018. Number: 5 Publisher: Nature Publishing Group.

[60] Bosiljka Tasic, Zizhen Yao, Lucas T. Graybuck, Kimberly A. Smith, Thuc Nghi Nguyen, Darren Bertagnolli, Jeff Goldy, Emma Garren, Michael N. Economo, Sarada Viswanathan, Osnat Penn, Trygve Bakken, Vilas Menon, Jeremy Miller, Olivia Fong, Karla E. Hirokawa, Kanan Lathia, Christine Rimorin, Michael Tieu, Rachael Larsen, Tamara Casper, Eliza Barkan, Matthew Kroll, Sheana Parry, Nadiya V. Shapovalova, Daniel Hirschstein, Julie Pendergraft, Heather A. Sullivan, Tae Kyung Kim, Aaron Szafer, Nick Dee, Peter Groblewski, Ian Wickersham, Ali Cetin, Julie A. Harris, Boaz P. Levi, Susan M. Sunkin, Linda Madisen, Tanya L. Daigle, Loren Looger, Amy Bernard, John Phillips, Ed Lein, Michael Hawrylycz, Karel Svoboda, Allan R. Jones, Christof Koch, and Hongkui Zeng. Shared and distinct transcriptomic cell types across neocortical areas. Nature, 563(7729):72–78, November 2018. Number: 7729 Publisher: Nature Publishing Group.

[61] Milad R. Vahid, Erin L. Brown, Chloé B. Steen, Wubing Zhang, Hyun Soo Jeon, Minji Kang, Andrew J. Gentles, and Aaron M. Newman. High-resolution alignment of single-cell and spatial transcriptomes with CytoSPACE. Nature Biotechnology, pages 1–6, March 2023. Publisher: Nature Publishing Group.

[62] Xiao Wang, William E. Allen, Matthew A. Wright, Emily L. Sylwestrak, Nikolay Samusik, Sam Vesuna, Kathryn Evans, Cindy Liu, Charu Ramakrishnan, Jia Liu, Garry P. Nolan, Felice-Alessio Bava, and Karl Deisseroth. Three-dimensional intact-tissue sequencing of single-cell transcriptional states. Science, 361(6400):eaat5691, July 2018. Publisher: American Association for the Advancement of Science.

[63] Yunguan Wang, Bing Song, Shidan Wang, Mingyi Chen, Yang Xie, Guanghua Xiao, Li Wang, and Tao Wang. Sprod for de-noising spatially resolved transcriptomics data based on position and image information. Nature Methods, 19(8):950–958, August 2022. Number: 8 Publisher: Nature Publishing Group.

[64] Runmin Wei, Siyuan He, Shanshan Bai, Emi Sei, Min Hu, Alastair Thompson, Ken Chen, Savitri Krishnamurthy, and Nicholas E. Navin. Spatial charting of single-cell transcriptomes in tissues. Nature Biotechnology, 40(8):1190–1199, August 2022. Number: 8 Publisher: Nature Publishing Group.

[65] Xiaoyu Wei, Sulei Fu, Hanbo Li, Yang Liu, Shuai Wang, Weimin Feng, Yunzhi Yang, Xiawei Liu, Yan-Yun Zeng, Mengnan Cheng, Yiwei Lai, Xiaojie Qiu, Liang Wu, Nannan Zhang, Yujia Jiang, Jiangshan Xu, Xiaoshan Su, Cheng Peng, Lei Han, Wilson Pak-Kin Lou, Chuanyu Liu, Yue Yuan, Kailong Ma, Tao Yang, Xiangyu Pan, Shang Gao, Ao Chen, Miguel A. Esteban, Huanming Yang, Jian Wang, Guangyi Fan, Longqi Liu, Liang Chen, Xun Xu, Ji-Feng Fei, and Ying Gu. Single-cell Stereo-seq reveals induced progenitor cells involved in axolotl brain regeneration. Science, 377(6610):eabp9444, September 2022. Publisher: American Association for the Advancement of Science.

[66] Joshua D. Welch, Velina Kozareva, Ashley Ferreira, Charles Vanderburg, Carly Martin, and Evan Z. Macosko. Single-Cell Multi-omic Integration Compares and Contrasts Features of Brain Cell Identity. Cell, 177(7):1873–1887.e17, June 2019.

[67] Håkan Wieslander, Philip J. Harrison, Gabriel Skogberg, Sonya Jackson, Markus Fridén, Johan Karlsson, Ola Spjuth, and Carolina Wählby. Deep Learning With Conformal Prediction for Hierarchical Analysis of Large-Scale Whole-Slide Tissue Images. IEEE Journal of Biomedical and Health Informatics, 25(2):371–380, February 2021. Conference Name: IEEE Journal of Biomedical and Health Informatics.

[68] Chenglong Xia, Jean Fan, George Emanuel, Junjie Hao, and Xiaowei Zhuang. Spatial transcriptome profiling by MERFISH reveals subcellular RNA compartmentalization and cell cycle-dependent gene expression. Proceedings of the National Academy of Sciences, 116(39):19490–19499, September 2019. Publisher: Proceedings of the National Academy of Sciences.

[69] Zizhen Yao, Cindy T. J. van Velthoven, Thuc Nghi Nguyen, Jeff Goldy, Adriana E. Sedeno-Cortes, Fahimeh Baftizadeh, Darren Bertagnolli, Tamara Casper, Megan Chiang, Kirsten Crichton, Song-Lin Ding, Olivia Fong, Emma Garren, Alexandra Glandon, Nathan W. Gouwens, James Gray, Lucas T. Graybuck, Michael J. Hawrylycz, Daniel Hirschstein, Matthew Kroll, Kanan Lathia, Changkyu Lee, Boaz Levi, Delissa McMillen, Stephanie Mok, Thanh Pham, Qingzhong Ren, Christine Rimorin, Nadiya Shapovalova, Josef Sulc, Susan M. Sunkin, Michael Tieu, Amy Torkelson, Herman Tung, Katelyn Ward, Nick Dee, Kimberly A. Smith, Bosiljka Tasic, and Hongkui Zeng. A taxonomy of transcriptomic cell types across the isocortex and hippocampal formation. Cell, 184(12):3222–3241.e26, June 2021.

[70] Chihao Zhang, Kangning Dong, Kazuyuki Aihara, Luonan Chen, and Shihua Zhang. STAMarker: Determining spatial domain-specific variable genes with saliency maps in deep learning, November 2022. Pages: 2022.11.07.515535 Section: New Results.

[71] Yan Zhou, Dong Yang, Qingcheng Yang, Xiaobin Lv, Wentao Huang, Zhenhua Zhou, Yaling Wang, Zhichang Zhang, Ting Yuan, Xiaomin Ding, Lina Tang, Jianjun Zhang, Junyi Yin, Yujing Huang, Wenxi Yu, Yonggang Wang, Chenliang Zhou, Yang Su, Aina He, Yuanjue Sun, Zan Shen, Binzhi Qian, Wei Meng, Jia Fei, Yang Yao, Xinghua Pan, Peizhan Chen, and Haiyan Hu. Single-cell RNA landscape of intratumoral heterogeneity and immunosuppressive microenvironment in advanced osteosarcoma. Nature Communications, 11(1):6322, December 2020.

[72] Jiaqiang Zhu, Lulu Shang, and Xiang Zhou. SRTsim: spatial pattern preserving simulations for spatially resolved transcriptomics. Genome Biology, 24(1):39, March 2023.

